# Basophils activate splenic B cells and Dendritic cells via IL-13 signaling in acute Traumatic Brain Injury

**DOI:** 10.1101/2025.05.20.655028

**Authors:** Florian Olde Heuvel, Jin Zhang, Fan Sun, Sruthi Sankari Krishnamurthy, Gizem Yartas, Marica Pagliarini, Michael K.E. Schäfer, Markus Huber-Lang, Francesco Roselli

**Author notes:** Correspondence: Prof. Dr. Francesco Roselli (MD, PhD) Centre for Biomedical Research Helmholtzstrasse 8 (R3.05)-89081 Ulm DE.

## Abstract

Peripheral consequences following traumatic brain injury (TBI) are characterized by both systemic inflammatory responses and autonomic dysregulation, with almost all peripheral organs affected. One of the main immune regulatory organs, the spleen, shows high interaction with the brain which is controlled by both circulating mediators as well as autonomic fibers targeting splenic immune cells. The brain-spleen axis does not function as a one-way street, it also shows reciprocal effects where the spleen affects neuroinflammatory and cognitive functions post injury. To date, systemic and splenic inflammatory responses are measured by cells or mediators located in circulation. Nevertheless, most of the signaling and inflammation post injury takes place in the organs. Therefore, we set out to investigate the early signaling landscape in the spleen following TBI, using phospho-proteomic signaling approaches and immunofluorescence stainings to investigate novel molecular and cellular players. Based on the signaling signature, we found a rapid influx of basophil granulocytes towards the spleen, which are recruited via CXCL1 expressed by B-cells and dendritic cells. The basophils activate B cells and dendritic cells (DCs) via the IL-13/IL-13Ra1 signaling pathway to enhance protein translation through the long non-coding RNA NORAD. The early recruitment of basophils and subsequent activation of B cells and DCs, is short lived and sets at 3dpi. Interestingly, the rapid recruitment of basophils is inhibited by ethanol intoxication in TBI. In conclusion, basophils recruitment to the spleen may serve as an early mediator of systemic inflammatory responses to TBI with potential implications for research on biomarkers and therapeutic targets.

## Introduction

Traumatic brain injury (TBI) has reported peripheral disturbances, characterised by both a systemic inflammatory response as well as an autonomic dysregulation in almost all distal organs[1,2]. TBI is associated with large scale immune modulatory effects showing both an increased systemic inflammation, by release of inflammatory cytokines and mediators[3–5], as well as suppression of immune responses, with a reduction in circulating immune cells, like lymphocytes and natural killer cells[6–9]. Interestingly, the TBI induced systemic inflammatory responses have been reported to last up to 60 days post injury[10], suggesting not only acute, but also long-term systemic inflammatory consequences. Therefore, the general systemic inflammatory response post TBI is highly relevant, since it may contribute to an enhanced vulnerability towards infections post injury.

One of the main immune regulatory organs, the spleen, is involved in primary innate immune responses such as antigen presentation followed by secondary adaptive immune response including the activation of lymphocytes and production of antibodies, as well as filtering of red blood cells[11]. Neurological conditions have been shown to modulate splenic immune functions, through a pathway known as the “brain-spleen axis”[12–14]. This interaction is modulated by both circulating mediators[15], as well as autonomic fibres targeting splenic immune cells[16,17]. In fact, stimulation of sympathetic and parasympathetic nerve fibers have been reported to inhibit inflammatory mediators, such as IL-6, IFN-y and TNF-a[18,19]. Interestingly, TBI has been associated with a reduction of adrenergic receptors in dendritic cells, which is associated with their increased maturation and TNF-a response[20]. Nevertheless, the brain-spleen axis upon TBI does not function as a one-way street; rather, it exhibits reciprocal effects as evidenced by alterations in the cognitive and neuroinflammatory responses following splenectomy post injury[21,22].

To date, most reports on the systemic, and in particular splenic, immune function upon injury have focused primarily on cellular effects and cytokines located in the systemic circulation, with less focus on in depth splenic molecular and cellular mechanisms. However, pharmacological interventions for systemic immune disturbances are largely targeting either molecular signaling pathways or specific cells within the inflamed tissue[23]. Together with the fact that immune responses occur within tissues rather than in the blood, we aimed at disentangling the early signaling responses in the spleen upon TBI.

In the present study we employed a phospho-proteomic signaling approach to identify a novel immunomodulatory mechanism in the acute phase upon TBI, involving the fast recruitment of basophils towards the spleen. Investigation of these basophils revealed a basophil to B-cell and Dendritic cell interaction, via IL-13 signaling. Followed by an enhanced protein translation, through a lncRNA NORAD fashion, in B-cells and DCs.

## Material and Methods

### Animals, traumatic brain injury model and ethanol treatment

This study is a post-hoc analysis of the spleen obtained from mice in previous studies[24–27]. This study was undertaken in accordance with the 3R principle, to reduce the number of animals in experimentation and increase the scientific output from sacrifice of animals. The experiments have been approved by the animal oversight committee at Ulm University and by the Regierungspräsidium in Tübingen (Reg. 1222). Wild-type (B6-SJL) male mice were bred locally, under standard housing conditions (24°C, 60%–80% humidity, 12/12 light/dark cycle, with *ad libitum* access to food and water). TBI was performed on mice p60-90, as previously reported[24–27]. Briefly, mice were administered buprenorphine (0.1 mg/kg, subcutaneously) followed by sevoflurane anaesthesia (2.5% in 97.5% O2). The skin was shaved and incised on the midline, after which the mice were positioned in the apparatus. The closed head weight drop was delivered by a weight of 333 g falling from a height of 2 cm, on the parietal bone. Mice were administered 100% O2, directly after the TBI, with monitoring of apnoea time. Control mice (sham) had the same treatments (analgesia, anaesthesia, skin incision and handling), without the TBI being administered. Ethanol treatment was performed as previously described[28]. Briefly, 100% synthesis grade ethanol was diluted in 0.9% NaCl to a final dilution of 32% volume/volume. Mice (20–25 g) were administered a volume of 400–500 µl of diluted ethanol (to obtain a concentration of 5 g/kg) by oral gavage 30 min before TBI. Four experimental groups were considered: saline administered, subjected to sham surgery (saline-sham, SS); saline administered, subjected to TBI (saline-TBI, ST); ethanol administered, subjected to sham surgery (ethanol-sham, ES); and ethanol administered, subjected to TBI (ethanol-TBI, ET).

### Tissue isolation

Three hours or three days post TBI, mice were sacrificed by cervical dislocation, followed by organ and blood harvesting for further processing. The spleen was dissected and snap frozen on dry ice. Tissue was either used for protein isolation for western blot or protein arrays, RNA isolation for RT-qPCR or tissue sectioning for immunofluorescence staining or RNAscope in situ hybridisation. Blood samples were collected in EDTA tubes and centrifuged at 5000g, 4°C, for 5 min, plasma was collected and frozen at -80°C for further use.

### Protein isolation and determination

Proteins were isolated from the spleen by adding RIPA buffer (150 mM NaCl, 50 mM Tris pH 7.6, 0.1% SDS, 1% Triton X-100, 1 mM EDTA), in a volume 4x the weight of the spleen (weight: volume), protease and phosphatase inhibitors (Roche tablets) were added in excess, followed by homogenization of the tissue using a tissue grinder. The homogenate was sonicated for 2 x 2 sec at 65%, followed by incubation at 4°C for 30 min. The homogenate was centrifuged for 20 min at 10.000g, the supernatant was moved to another tube and the pellet was discarded. The protein concentration was determined using the BCA assay (Thermo Fischer) according to the manufacturer’s instructions. Briefly, the homogenate was diluted 20x in RIPA buffer and added to the wells together with the standard in duplicate, 200ul of working reagent (50:1, reagent A:B) was added to each well and incubated at 37°C for 30 min. The absorbance was measured at 562 nm with a plate reader. Sample concentration was determined by calculation from the standard curve.

### SDS-page and Western blot

50 µg of protein with 1x loading buffer and 5% beta-mercaptoethanol was heated to 95°C for 5 min, after which it was loaded and run on 8-12% SDS-PAGE gel at 100V until separation of the proteins (1-2h approx.). Then proteins were transferred to a nitrocellulose membrane at 100V for 1.5-2h, followed by blocking (5% BSA, 1% TBS-T) for 1h at RT while shaking. Membranes were incubated with primary antibodies (supplementary table 1), diluted in blocking buffer, at 4°C overnight while shaking. The membranes were washed with TBS-T 3 x 10 min at RT while shaking, followed by incubation of the HRP secondary antibodies, diluted in blocking buffer, for 1h at RT on the shaker. A last wash was performed with TBS-T 3 x 10 min at RT while shaking, followed by detection by adding enhanced chemiluminescence (ECL) on the membrane and imaging using a Las4000 ImageQuant (GE healthcare). Analysis was performed with ImageJ, measuring the mean gray value of each band and normalised to the loading control.

### Antibody array assay

AKT signaling antibody array kit was performed according to manufacturer’s instructions (Pathscan AKT signaling antibody array kit; CST). Briefly, slides were blocked by adding 75 µl of blocking buffer to each well for 15 min at RT, followed by 100 µl, 1 mg/ml lysate incubation at 4°C overnight while shaking. Slides were washed 4 x 5 min using washing buffer, followed by an incubation of 150 µl detection antibody cocktail to each well for 1h at RT while shaking. Slides were washed 4 x 5 min with washing buffer, followed by an incubation with Cy5-streptavidin (Thermo Fischer; 1:1000) for 20 min at RT, protected from light. Slides were washed a final time 4 x 5 min with washing buffer and air dried. The slides were scanned using a GenePix 4000B array scanner (Molecular Devices, LLC). Spotting of the arrays was performed semi-automatically with manual correction of each spot, followed by extraction of the data using the GenePix pro 6.1 software.

Cytokine and vascular factors (angiogenesis) antibody array kits were performed according to manufacturer’s instructions (Proteome Profiler Mouse XL Cytokine Array and Proteome Profiler Mouse Angiogenesis Array Kit; R&D systems). Briefly, membranes were blocked for 1h at RT in buffer 6 while shaking, followed by an overnight incubation at 4°C of 200ug proteins (diluted in buffer 4 to a total volume of 1.5ml). Membranes were washed 3 x 10 min in 1x wash buffer at RT while shaking, followed by an incubation of detection antibody (diluted in buffer 4 to a total volume of 1.5ml) at RT for 1h while shaking. The membranes were washed 3 x 10 min in 1x wash buffer at RT while shaking, followed by incubation of streptavidin-HRP for 30 min at RT while shaking. The membranes were washed a final time 3x 10 min in 1x wash buffer at RT while shaking, followed by incubation of chemi reagent mixture for 1 min. Detection was performed using a chemidoc system (BioRad). Analysis was performed with ImageJ, measuring the mean gray value of each spot, background subtracted and normalised to the positive control.

Phospho-explorer antibody array was performed according to manufacturer’s instructions (Phospho Explorer Antibody Array; Fullmoon bio). Briefly, 100ug of Protein samples, diluted with labeling buffer to a total volume of 75ul, were biotinylated by adding 3ul of a biotin/DMF solution and incubated for 2h at RT, with vortexing every 10 min. Biotinylation was stopped by adding 35ul of stopping reagent and incubated for 30 min at RT, with vortexing every 10 min. Glass arrays were taken from 4°C, left in the package, and allowed to warm up to RT for 1h. The arrays were removed from the package and left at RT for another 30 min. The arrays were blocked for 45 min at RT on the shaker at 55 rpm. After blocking, the arrays were washed with Milli-Q grade water ten times. The biotinylated sample was combined with 6 ml Coupling Solution. The arrays were incubated with protein coupling mix on the orbital shaker rotating at 35 rpm for 2 h at RT. After coupling, the arrays were washed with 1x Washing Solution on the shaker rotating at 55 rpm for 3 x 10 min. The arrays were extensively rinsed with Milli-Q water as before. The arrays were submerged in the Detection Buffer with 0.5 mg/ml Cy5-streptavidin on the shaker rotating at 35 rpm for 20 min at RT in the dark. The arrays were washed by Washing Solution and rinsed with Milli-Q water as before. After drying with compressed nitrogen, the slides were scanned using a GenePix 4000B array scanner (Molecular Devices, LLC). Spotting of the arrays was performed semi-automatically with manual correction of each spot, followed by extraction of the data using the GenePix pro 6.1 software.

### ELISA assay

The ELISA assays for CXCL1, CXCL13, CCL6 and IL-5 on spleen and plasma were performed according to manufacturer’s instructions (mouse DuoSet ELISA; R&D systems). Briefly, the plate was incubated overnight at RT with 100 µl of capture antibody solution, followed by 3 x washing step with wash buffer. The plate was blocked for 1h at RT with 300 µl of reagent diluent, followed by 3 x washing step with wash buffer. 100 µl of standards and samples were added in various dilutions in reagent diluent (standards undiluted; CCL6: spleen 1:100, plasma 1:2; CXCL1 spleen 1:20, plasma 1:2; CXCL13 spleen 1:100, plasma undiluted; IL-5: spleen 1:25, plasma undiluted) and incubated for 2h at RT, followed by 3 x washing step with wash buffer. 100 µl of detection antibody was added and incubated for 2h at RT, followed by 3 x washing step with wash buffer. 100 µl of streptavidin-HRP was added and incubated for 20 min at RT, followed by 3 x washing step with wash buffer. 100 µl of substrate solution was added and incubated for 20 min at RT (covered from light), followed by 50 µl of stop solution to each well. The optical density was determined using a microplate reader at 450 nm (with a wavelength correction at 540 nm). Data was extracted and concentrations were calculated against a 4-PL curve (4-parameter logistics curve), followed by corrections of dilutions.

### RNA isolation and quantitative RT-PCR

RNA was isolated from the spleen using QIAzol (Qiagen, Germany) by disrupting and homogenizing the tissue in 1 ml QIAzol, 250 µl of chloroform was added and vortexed for 15 s. The samples were placed at RT for 10 min and centrifuged for 10 min 12,000×*g* at 4°C to achieve phase separation. The top layer (containing RNA) was moved to another tube and precipitated with a similar amount of isopropanol. The samples were placed at RT for 10 min and centrifuged for 10 min 12,000×*g* at 4°C, with visible pellet formation. Isopropanol was discarded and 1 ml of 75% ethanol was added and mixed, to wash the pellet. The samples were centrifuged for 10 min 8,000×*g* at 4°C, ethanol was discarded, and the samples were air dried. The RNA pellet was redissolved in 20 µl RNAse-free dH_2_O. RNA concentration was determined by NanoDrop. Reverse transcription was performed by adding 5 µl random hexamers (Biomers, Germany) to 1 μg RNA (total volume 40 µl diluted in dH_2_O) and incubated for 10 min at 70°C. The samples were placed on ice and a master mix of 0.5µl reverse transcriptase (Promega, Germany), 0.5 µl RNase Inhibitor (RiboLock, Thermo Scientific, Germany), 2 µl dNTPs (Genaxxon, Germany), and 12 µl reverse transcriptase buffer (Promega, Germany). The samples were incubated for 10 min at RT and placed in a water bath for 45 min at 42°C. The samples were incubated for 3 min at 99°C, placed on ice, and frozen until further use.

qPCR was performed on the Quantstudio 1 real-time PCR machine (Thermo fischer) with the Power SYBR-green PCR master mix (Applied biosystems). Two microliters of sample cDNA were used in a total volume of 10 µl (3 µl primer mix and 5 µl of Sybr-green) in a 96-well plate, all samples were duplicated, and the housekeeping gene Actin was used as a control. The Ct values obtained from the lightcycler were normalized according to the following equation: 2−ΔΔCt to obtain the fold change.

### Tissue sectioning and immunofluorescence staining

Frozen spleen tissue was embedded in OCT (Tissue-Tek, The Netherlands), 10 μm sections were cut using the cryostat and mounted on superfrost plus glass slides. The slides were stored for at least 24h at −80°C and washed for 5 min in 1 × PBS, followed by a 15-min fixation step of the sections in 4% PFA at 4°C. blocking of the sections was done in blocking buffer (3% BSA, 0.3% Triton X-100, PBS) for 2 h at RT. The primary antibodies (supplementary table 1) were diluted in blocking buffer and incubated overnight at 4°C, followed by 3 × 30 min washes in PBS at RT. The secondary antibodies were diluted in blocking buffer and incubated for 2 h at RT, followed by 3 × 30 min washes in PBS. The sections were mounted using Fluorogold prolong antifade mounting medium (Invitrogen, Germany).

### Single mRNA in situ hybridisation

mRNA in situ hybridisation was performed according to manufacturer’s instructions (ACDBio, RNAscope, Newark, CA, USA fluorescence *in-situ* hybridization for fresh frozen tissue, all reagents and buffers were provided by ACDBio[29]. Briefly, frozen spleen tissue was embedded in OCT (Tissue-Tek, The Netherlands), 10 μm sections were cut using the cryostat and mounted on superfrost plus glass slides. The slides were stored for at least 24 h at -80°C and washed for 5 min in 1 x PBS, followed by a 15 min fixation step in 4% PFA at 4°C. The sections were dehydrated in a series of ascending ethanol concentrations (50% - 70% - 100%) for 5 min each, followed by a last time in 100% ethanol for 5 min. Sections were air dried and covered in 30% hydrogen peroxide for 10 min at RT, followed by 2x wash in PBS by moving the sections up and down 5x. Sections were covered in protease IV and incubated at 40°C for 30 min, followed by 2 x 2 min washing in PBS. The probes (CXCL1 and NORAD) were added and incubated at 40°C for 2h, followed by 2 x 2 min washing in wash buffer. Then, amplification 1 was added to the sections and incubated at 40°C for 30 min followed by 2 × 2 min washing step with wash buffer. Next, amplification 2 was added to the sections and incubated at 40°C for 30 min followed by 2 × 2 min washing step with wash buffer. As a final amplification step, amplification 3 was added and incubated at 40°C for 15 min, followed by 2 × 2 min washing step with wash buffer. HRP corresponding to the channel was added to the sections and incubated at 40°C for 15 min, followed by 2 x 2 min washing step with wash buffer. The diluted fluorophore (1:1500 for CXCL1 and 1:1000 for NORAD; in TSA buffer) was added to the sections and incubated at 40°C for 30 min, followed by 2 x 2 min washing step with wash buffer. The remaining HRP was blocked by adding HRP-blocker to the sections and incubated at 40°C for 15 min, followed by 2 x 2 min washing step with wash buffer. Sections were blocked using blocking buffer (3% BSA, 0.3% Triton X-100, PBS) at RT for 1h, followed by primary antibody, diluted in blocking buffer, incubation at 40°C overnight. The sections were washed 3 x with PBS-T (1% triton-x100) at RT for 20 min each, followed by secondary antibody incubation (diluted in blocking buffer) at RT for 2h. The sections were washed 3 x with PBS-T at RT for 20 min each, and mounted using Fluorogold prolong antifade mounting medium (Invitrogen, Germany).

### Image acquisition

Confocal images were acquired with a laser-scanning confocal microscope (LSM Leica DMi8), with a 20x oil (NA 0.75) and 40x oil (NA 1.3) objective in a 1024 × 1024 pixel resolution and a 12-bit format. A z-stack of 7-8 optical sections spanning 14-16 μm (step size of 2 μm) and 1.5 optical zoom was made. A tilescan of 2 x 2 was made spanning the entire follicle (with 20x objective), or a part of the follicle (with the 40x objective). Imaging parameters were set to obtain signals from the stained antibody/probe while avoiding saturation. All fluorescent channels were acquired independently to avoid fluorescence cross-bleed. For each treatment, three spleen sections per animal were randomly selected and imaged.

### Super-resolution STED imaging

Super-resolution STED imaging was performed on a Stedycon module (Abberior instruments, Germany) fixed to a Zeiss microscope fitted with a 100× (NA 1.4) oil objective. Images were acquired in both confocal and STED modes; imaging parameters were set to obtain signals from the staining while avoiding saturation. For the imaging, Abberior Star 580 and Abberior Star 635 dyes were used, with an excitation laser of 561 nm for the Star 580 and 640 nm for the Star 635 and a STED laser of 775 nm for both channels. The pixel size was set to 25 nm, with a pixel dwell time of 10 µs and line accumulation of 6 µs. A positive basophil was selected, and a single optical section was acquired in correspondence with the optical plane.

### Image analysis

Image analysis was performed using the ImageJ software. Images were background subtracted and smoothed. For the immunofluorescent staining, images were contrasted to assess positive and negative cells, positive cells were counted manually using the cell counter plugin and cells were depicted as either absolute number per area or as percentage of cell type. For the in-situ hybridisation, images were contrasted to assess positive and negative cells, cells positive for the mRNA marker and cellular marker were counted manually using the cell counter plugin and values were depicted as percentage of cell type. For the proximity analysis of pIL-13ra1+ close to IL-13+ cells, 4 fixed size circles (40, 60, 80 and 100 micron) were placed around the IL-13+ cell and the amount of pIL-13Ra1+ cells were manually counted in each circle using the cell counter plugin. For the proximity analysis of NORAD+ cells close to MCPT8+ cells, a fixed size circle (40 micron) was placed around the MCPT8+ cell and the ratio of NORAD+ and either CD19+ or CD11c+ were manually counted using the cell counter plugin. For the Super-resolution STED images, deconvolution of all images was performed using the web based TRUESHARP image boosting tool from Abberior star[30], with following parameters; Number of iterations: 5, PSF size: 2, background size: 16 px, background weight: 1. For the IL-13/pIL-13ra1 analysis, an intensity plot was traced along an IL-13 cluster (in positive basophils) towards the pIL-13Ra1 cluster; the distance from IL-13 to pIL13-Ra1 was computed. For the Actin/pIL-13Ra1 analysis an intensity plot was traced over the actin of basophil towards the pIL-13ra1 positive cell, the distance from the actin to the pIL-13ra1 was computed.

### Antibody array analysis and bio-informatics

Antibody array analysis was performed using the R-studio version 4.3.1, using the limma package, utilizing linear models to model for experimental designs. Normalization within and between arrays was done using the cyclicloess method, to remove intensity-dependent biases between different samples and thereby making them more comparable. Then the arrays were corrected for any batch effects using the ComBat function, which estimates the mean and variance for each batch and adjusts the data accordingly. Then we proceeded towards differential expression, employing empirical Bayes methods that samples information across all proteins on the array, moderating the standard errors and improving the power to detect differential expression. The output was obtained using the topTable function to extract lists of differentially expressed proteins and their associated statistics like log fold-change, p-values, adjusted p-values, etc. as in Supplementary file 1.

Bio-informatic analysis was performed using the online “Immgen MyGeneset” database (Immunological Genome project; https://www.immgen.org/)[31]. The significant targets were imported into the database, the cell type selected and the analysis run using the Immgen ULI RNA-seq database. The heatmap and W-plot were extracted with the median normalised function showing the involvement of immune cell types dependent on the significant targets. The KEGG pathway analysis was performed using the online ShinyGo database[32], using a FDR of 0,05 with a min and max pathway size of 2 and 5000 respectively. The significant KEGG pathways were extracted and preselected based on immunological function.

### Statistical analysis

Data analysis was performed using the Graphpad prism version 8 software or R software suite Version 4.3.1. Statistical analysis was performed using animals as a biological unit; whenever appropriate, single data points and averages per animal are depicted in the graphs. All datasets were tested for normality using the Shapiro-Wilks test. When comparing two groups, an unpaired t-test or Mann-Whitney test was used. When comparing multiple groups, ANOVA with Sidak’s multiple correction or Kruskal-Wallis test with Dunn’s multiple correction were used, depending on normality.

## Results

### Splenic large-scale signaling landscape following TBI points toward basophil influx

Since immune reactions largely take place within tissues rather than blood, we set out to investigate the early immunoregulatory events taking place after TBI by focusing on the spleen. As a first step, we verified that TBI induced acute changes (3h post injury) in the splenic signaling landscape. We probed spleen extracts obtained 3h after TBI (or sham surgery) with a 16-target glass based fluorescent antibody array, focused on Erk/AKT/mTOR signaling. We identified significant differences in 7 out of 16 targets (Figure 1A), including increased phosphorylation of ERK1/2 (thr202/204), Bad (ser112) and GSK-30 (ser21), as well as decreased phosphorylation of Akt (ser473; thr308), RSK1 (ser380) and PDK1 (ser241). The upregulation of ERK phosphorylation (Figure 1B, C) and of S6-RP phosphorylation (pS6, S235/236 - a marker for increased translation; Figure 1B, D) was confirmed by WB, which demonstrated the significant activation of early signaling and cellular responses in the spleen upon TBI (pS6: p = 0.0367; pERK1/2: p = 0.0282; Figure 1B-D).

**Figure 1:**
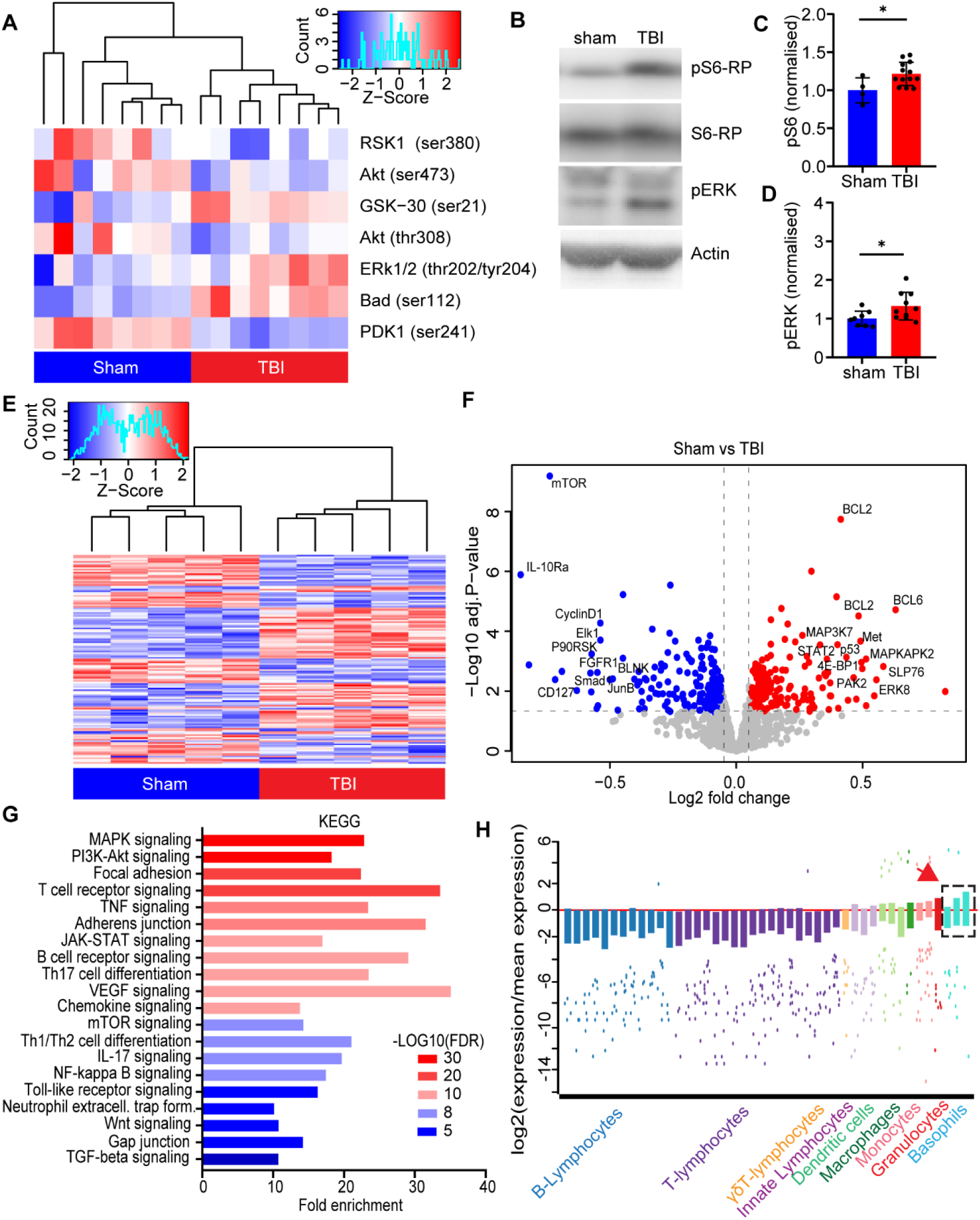
TBI results in large-scale signaling resulting in a basophils influx to the spleen. **A.** The heatmap shows significantly altered phosphorylation targets (down: RSK1, AKT and PDK1; up: GSK-30, ERK1/2 and BAD) from the AKT signaling pathway in mouse spleen 3h post sham or TBI treatment. N = 8. **B-D.** Western blot analysis in spleen samples reveals increased phosphorylation of S6 ribosomal protein (Sham vs TBI: 1.0±0.2 vs 1.3±0.4; p = 0.0367; t = 2.279) and ERK1/2 (Sham vs TBI: 1.0±0.2 vs 1.2±0.2; p = 0.0282; t = 2.428) 3h post TBI. N = 4-13. **E-F.** Heatmap and volcanoplot, from large scale phospho-signaling array revealed 135 (out of 1184) targets altered in the spleen post TBI. Red = up-phosphorylated, blue = down-phosphorylated, several targets marked in the volcanoplot. N = 5. **G.** KEGG (Kyoto Encyclopedia of Genes and Genomes) analysis of phospho-proteomic targets linked to immune related function and biological processes. **H.** W-plot, extracted from the ImmGen My Geneset database, showing selected immune cells on the x-axis and the log2 expression of the significant targets from the phospho-proteomic dataset. The analysis shows a strong involvement of basophils (red arrow) in the spleen post TBI. Data is shown as mean±SD, z-scores for the heatmaps or fold enrichment with log10(FDR) for the KEGG analysis. *: *p* < 0.05. A complete list of altered phosphorylated targets can be found in supplementary information 1.

Next, we characterized in depth the signaling events unfolding in the spleen upon TBI. We performed a broad phospho-proteomic analysis using phospho-explorer arrays with 1318 targets from over 30 signaling pathways. We identified 135 significant differential phosphorylation events occurring upon TBI (Figure 1E-F), with several notable targets having increased phosphorylation involved in an inflammatory response including STAT2, MAP3K7 and SLP76[33–35] or a decreased phosphorylation of targets involved in depression or homeostasis of inflammation, such as; IL-10Ra1, P53 and SMAD1[36–38]. Interestingly, several targets related to increased metabolic signaling, like 4E-BP1 and ERK showed an increase in phosphorylation, as well as mTOR that revealed a decreased phosphorylation at site Thr2446, which has been associated with decrease in nutrients and is inversely related to S6 phosphorylation[39]. Highlighting the multifaceted splenic signaling architecture triggered by TBI. KEGG pathway analysis revealed a significant enrichment of pathways involved in an increased inflammatory response like; T-cell receptor signaling, TNF signaling, JAK-STAT signaling, B-cell receptor and Toll-like receptor signaling, as well as an involvement of chemotaxis and cell-to-cell contact like; focal adhesion, adherens junctions, chemokine signaling and gap junctions (Figure 1G).

To identify the immune cells subpopulations involved in the signaling events, we exploited the deconvolution algorithm provided by the Immunological Genome Project (available at: https://rstats.immgen.org/MyGeneSet_New/index.htm[31]). When provided with the 135 proteins displaying altered phosphorylation, the algorithm indicated the ongoing regulation of DCs, macrophages and granulocytes, with a specifically strong involvement of basophils (Figure 1H, arrow).

Taken together, the phosphoproteomic data reveals a rapid and sustained activation of signaling in the spleen upon TBI, in particular involving DCs, granulocytes and, surprisingly, a substantial involvement of basophils in the spleen.

### TBI results in an increased chemoattraction of basophils in splenic follicles

Our initial screenings indicated a possible involvement of basophils in the early systemic inflammatory response to TBI. We sought direct, spatial confirmation of increased basophil influx by immunofluorescent staining of thin spleen sections with the pan-granulocyte marker Ly6G, together with a more specific basophil marker CCR3[40–42] and CD19 as a B-cell marker (to provide architectural context for germinal center and marginal zone of follicles; Figure 2A).

**Figure 2:**
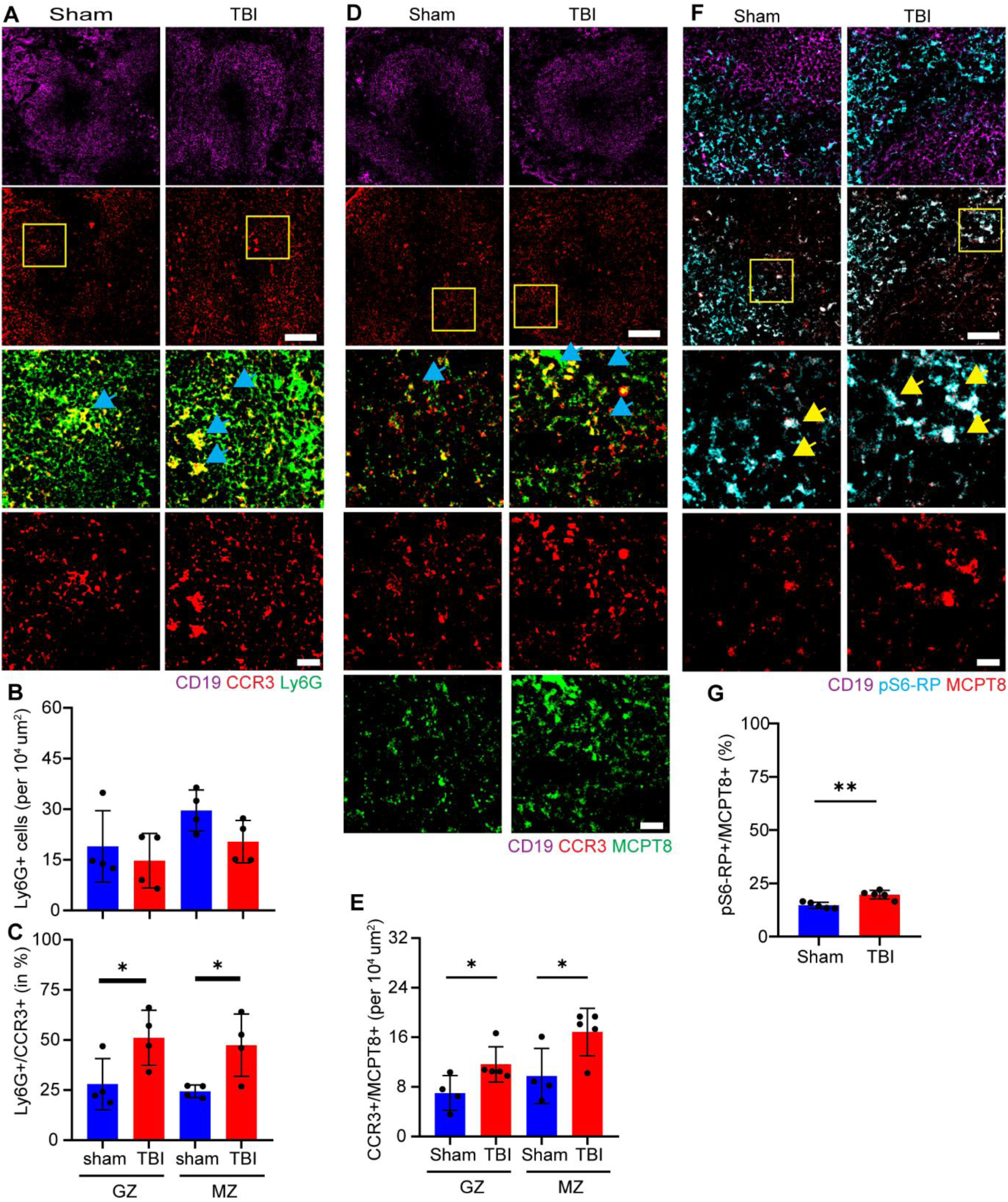
TBI induces chemoattraction of basophils in splenic follicles. **A-C.** Immunofluorescence staining of spleen sections with CD19, Ly6G and CCR3 show no differences in the number of Ly6G+ cells 3h post TBI. However, a significant increase was observed in Ly6G+/CCR3+ cells in the germinal center (Sham vs TBI: 27.9±12.8% vs 51.1±13.8%; p = 0.0494; t = 2.456) and marginal zone (Sham vs TBI: 24.4±3.2% vs 47.4±15.6%; p = 0.0275; t = 2.895) of splenic follicles. N = 4, scale bar overview: 100 µm; scale bar insert: 20 µm. **D, E.** Immunofluorescence staining of spleen sections with CD19, CCR3 and MCPT8 show a significant increase in CCR3+/MCPT8+ cells in the germinal center (Sham vs TBI: 7±2.8% vs 11.6±2.8%; p = 0.0451; t = 2436) and marginal zone (Sham vs TBI:9.7±4.5% vs 16.9±3.8%; p = 0.0361; t = 2.8586) of splenic follicles. N = 4, scale bar overview: 100 µm; scale bar insert: 20 µm. **F, G.** Immunofluorescent staining of spleen sections with CD19, pS6-RP and MCPT8 show a significant increase in pS6+/MCPT8+ cells (Sham vs TBI: 14.7±1.5% vs 19.7±2.0%; p = 0.002; t = 4.520) in splenic follicles. N = 5, scale bar overview: 50 µm; scale bar insert: 10 µm. Data is shown as mean±SD. *: *p* < 0.05, **: p<0.01.

Ly6G immunoreactivity was highly enriched mainly around and inside the follicles. The density of Ly6G positive cells was not altered by TBI, showing approx 20-25 positive cells per 10^4^ µm^2^ in both the germinal center and marginal zone (with a slightly larger number in the marginal zone; Figure 2A-B). On the other hand, the number of Ly6G+/CCR3+ cells showed a significant increase upon TBI in both the germinal center and marginal zone of the follicles (germinal center: p = 0.0494; marginal zone: p = 0.0275; Figure 2A, C).

In an independent experiment, an alternative basophil marker, MCPT8[43,44], was used, together with CCR3 and CD19 (Figure 2D). TBI caused a significant increase in the ratio CCR3+/MCPT8+ cells in both the germinal center as well as the marginal zone (Germinal center: p = 0.0451; marginal zone: p = 0.036; Figure 2D-E), suggesting that TBI enhances the chemoattraction of basophils into splenic follicles. In order to investigate if the basophils directly contributed to the increased activity and signaling post TBI, we immunolabelled spleen sections with antibodies against ribosomal pS6, together with MCPT8 and CD19 for cell-type identification. TBI resulted in a significant increase in the ratio of MCPT8+/pS6+ cells in splenic follicles (p = 0.002; Figure 2F-G), suggesting an increase in protein synthesis rates of basophils upon TBI.

This data indicated not only an increased chemoattraction of basophils into the spleen post injury, but also their enhanced translational activity in splenic follicles.

### Splenic B-cells and Dendritic Cells attract Basophils, via chemokine CXCL1, upon TBI

The migration, chemotaxis and activation of immune cells towards the organ of interest occurs through distinct cytokine and chemokine signals[45,46]. Several cytokines, chemokines and immune mediators have been found to regulate basophil chemotaxis and activation[47–49]. To identify immune mediators involved in TBI-induced chemotaxis and homing of basophils in the spleen, we employed two distinct cytokine antibody arrays, one focused on chemokines and immune mediators (111 targets) and a second one focused on vascular activation factors (40 targets).

Principal component analysis (PCA) shows distinct clustering of the sham and TBI groups (Figure 3A, C). The differential expression analysis revealed the upregulation of 14 proteins and downregulation of 10 proteins, upon TBI. Several of upregulated proteins are chemokines or mediators involved in chemotaxis, such as CXCL1, CXCL13, CCL6, MCP-1 (CCL2), fractalkine (CX3CL1) and GM-CSF (Figure 3B, D; red arrows), in fact upregulation of CXCL1 was confirmed in both assays. Notably, this protein has been associated in the recruitment or activation of basophils[50]. Interestingly, IL-5, a critical regulator of the activation of basophils[51], shows a downregulation (Figure 3D; blue arrow).

**Figure 3:**
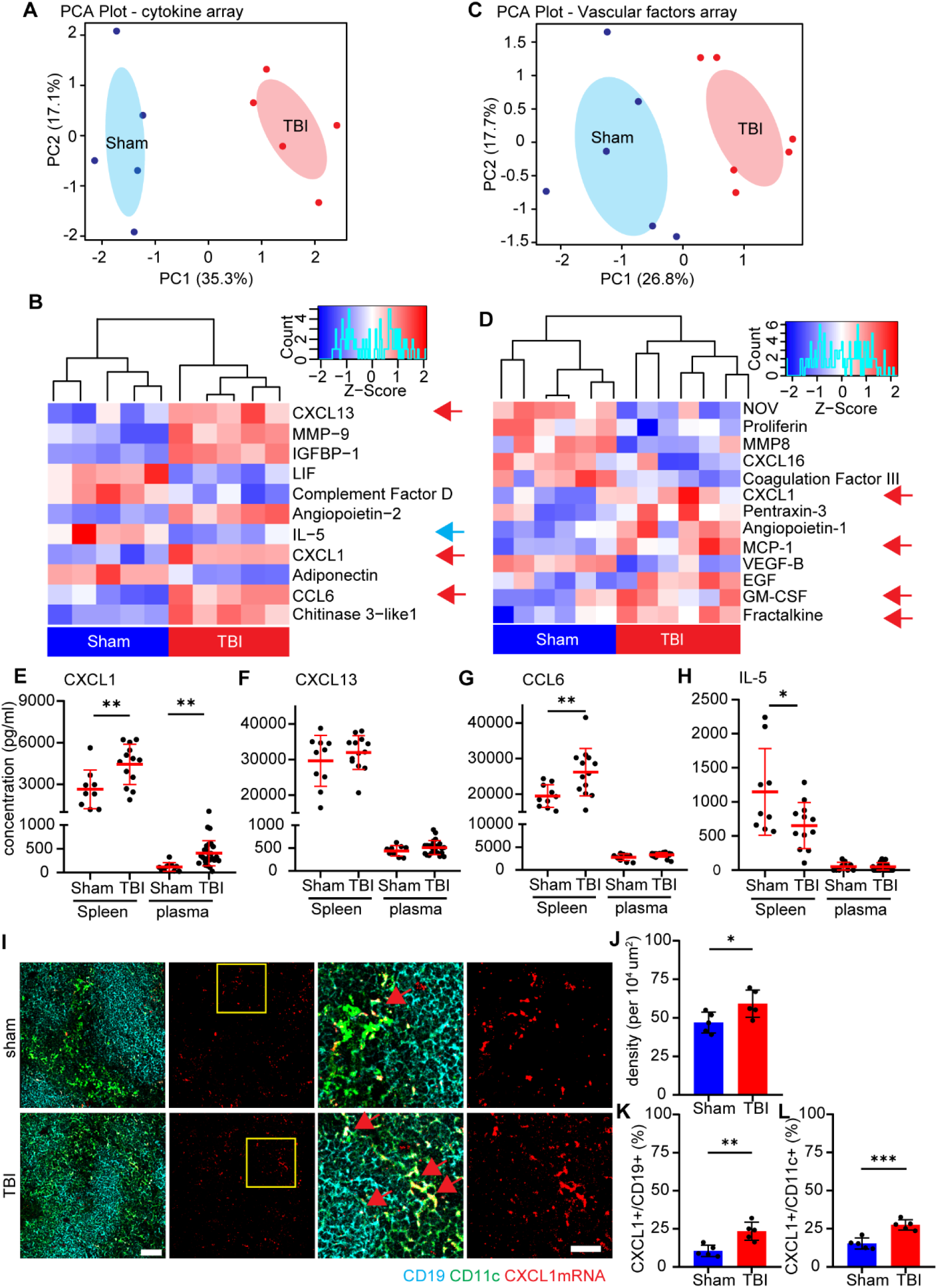
Chemoattraction of basophils upon TBI occurs through CXCL1 expression in B-cells and DCs. **A.** Principal component analysis (PCA) of the cytokine array revealed distinct clustering of the sham and TBI treatment groups. N = 5. **B.** The heatmap of the cytokine array showed the significantly altered cytokines/chemokines in the spleen 3h after TBI. N = 5. **C.** Principal component analysis (PCA) of the vascular factors array revealed distinct clustering of the sham and TBI treatment groups. N = 6. **D.** The heatmap of the vascular factors array showed the significantly altered cytokines/chemokines in the spleen 3h after TBI. N = 6. **E-H.** ELISA assays verification, in a larger subset of samples, showed the significant upregulation of CXCL1 (Sham vs TBI: 2642±1403 pg/ml vs 4439±1451 pg/ml; p = 0.0090; t = 2.893) and CCL6 (Sham vs TBI: 19466±3209 pg/ml vs 26175±6681 pg/ml; p = 0.0083; t = 2.916), a significant downregulation of IL-5 (Sham vs TBI: 1146±637 pg/ml vs 654±337 pg/ml; p = 0.0331; t = 2.298) and no difference in CXCL13 in the spleen. Plasma samples showed the significant upregulation only in CXCL1 (Sham vs TBI: 116±92 pg/ml vs 408±264 pg/ml; p = 0.0018; t = 3.387), with no differences observed in CXCL13, CCL6 and IL-5. Sham N = 10; TBI N = 13. **I-L.** Single mRNA RNAscope revealed the significant increase of CXCL1 mRNA puncta in the whole spleen (Sham vs TBI: 47±7 vs 59±9; p = 0.0377; t = 2.488), as well as an increase of CXCL1+/CD19+ (Sham vs TBI: 11±4% vs 23±6%; p = 0.0033; t = 4.125) and CXCL1+/CD11c+ (Sham vs TBI: 15±4% vs 28±3%; p = 0.0006; t = 5.484) cells after TBI. N = 5. scale bar overview: 50 µm; scale bar insert: 10 µm. Data is shown as mean±SD. *: *p* < 0.05, **: p<0.01, ***: p<0.001.

We expanded the investigation of a subset of mediators, namely CXCL1, CXCL13, CCL6 and IL-5, by performing ELISA quantification in both spleen protein extracts and plasma in a larger cohort of TBI samples. The assay independently verified the significant increase of CXCL1 (p = 0.0090; Figure 3E), CCL6 (p = 0.0083; Figure 3F) and decrease of CCL5 (p = 0.0331; Figure 3H) in the spleen, but did not confirm the upregulation of CXCL13 (Figure 3G). Interestingly, when plasma was considered, among the four mediators, only CXCL1 displayed a significant upregulation (p = 0.0018; Figure 3E); notably, CXCL1 absolute levels were significantly lower in the plasma than in the spleen, indicating a gradient leading migration from the blood to the spleen.

Since B-cells, dendritic cells and CD4+ T cells have been reported to express CXCL1 in various disease settings[52–54], we performed single mRNA in situ hybridisation with a co-immunostaining for CD19, CD11c and CD4, to identify the cellular source responsible of CXCL1 upregulation. Analysis of in situ hybridisation experiments revealed an overall significant increase of CXCL1 mRNA in the spleen upon TBI (p = 0.0377; Figure 3I, J), predominantly localized to CD19+ and CD11c+ cells (Figure 3I) and almost absent expression in CD4+ cells (Suppl. Figure 2). Indeed, TBI resulted in a significant increase in the fraction of double-positive CXCL1+/CD19+ cells and CXCL1+/CD11c+ cells (CD19: p = 0.0033; CD11c: p = 0.0006; Figure 3I, K, L) but no change in CD4+/CXCL1+ cells (Suppl. Figure 2).

These data suggest that TBI induces the expression of chemoattractant CXCL1 in CD19+ B cells as well as CD11c+ DCs, but not CD4+ T cells, which attracts basophils towards the spleen.

### Basophils activate B-cells and DCs via IL-13/IL-13Ra1 signaling upon TBI

Basophils are known to contribute to immunomodulation through their high level of IL-13 expression, which in turn promotes a Th2 adaptive immunity response[55,56]. Thus, we hypothesized that the basophils influx in the spleen may strongly affect the local IL-13 signaling. We performed an immunofluorescence staining with MCPT8 and IL-13 in thin spleen sections of sham and TBI treated mice. We showed a significant increase of MCPT8+ cells (p = 0.0256; Figure 4A, B) as well as a significant increase of IL-13+/MCPT8+ cells upon TBI (from 50% to 80%; p<0.05; Figure 4A, C).

**Figure 4:**
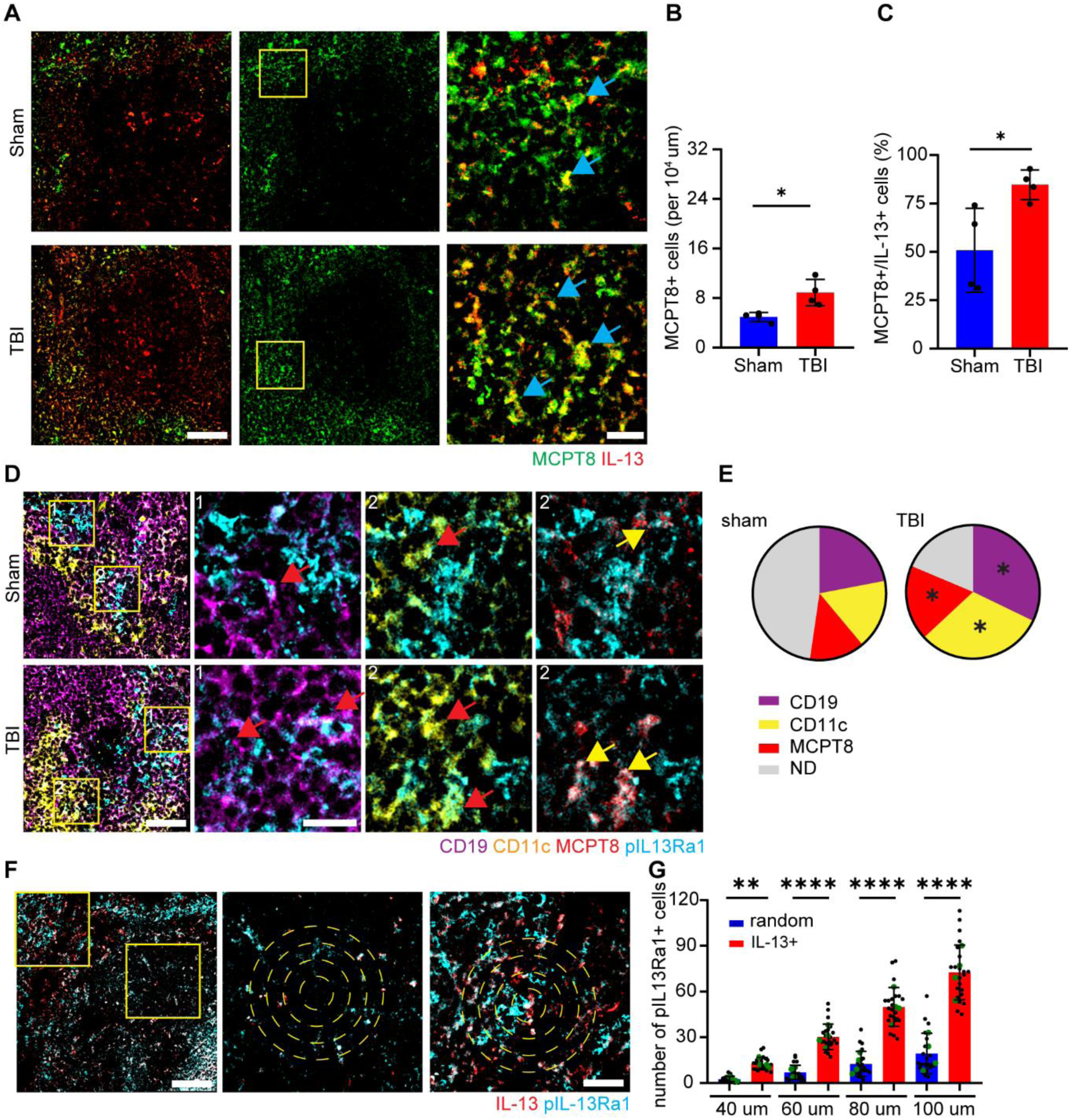
Basophils activate B-cells and DCs via IL-13 signaling post TBI. **A-C.** Immunofluorescence staining of spleen sections with MCPT8 and IL-13 revealed the significant increase in MCPT8+ cells per area (Sham vs TBI: 5±1 vs 9±2; p = 0.0129; t = 3.495) and the significant increase of MCPT8+/IL-13+ cells post TBI (Sham vs TBI: 50.8±21.7% vs 84.7±7.6%; p = 0.0256; t = 2.949). N = 4, scale bar overview: 100 µm; scale bar insert: 50 µm. **D, E**. Immunofluorescence staining of spleen sections with CD19, CD11c, MCPT8 and pIL-13Ra1 showed the significant increase in pIL-13Ra1+/CD19+ (Sham vs TBI: 22±4% vs 32±6%; p = 0.0305; t = 2.817), pIL-13Ra1+/CD11c+ (Sham vs TBI: 17±4% vs 31±10%; p = 0.0465; t = 2.501) and pIL-13Ra1+/MCPT8+ (Sham vs TBI: 13±2% vs 18±2%; p = 0.0176; t = 3.244) cells post TBI. N = 4, scale bar overview: 50 µm; scale bar insert: 20 µm. **F, G.** Immunofluorescence staining of thin spleen sections with IL-13 and pIL-13Ra1 revealed a significant increase of pIL-13Ra1+ cells in close proximity to IL-13+ cells as compared to a random location (IL-13+ vs random; 40 µm: 3±2 vs 13±4; 60 µm: 7±5 vs 30±8; 80 µm: 12±8 vs 50±13; 100 µm: 19±13 vs 73±18; p = 0.0029; p < 0.0001; p < 0.0001; p < 0.0001; F_(7,192)_ = 138.1). N = 4, scale bar overview: 100 µm; scale bar insert: 50 µm. Data is shown as pie-charts or mean±SD. *: *p* < 0.05, **: p<0.01, ***: p<0.001.

IL-13 signals through the type 1 receptor, consisting of IL-13Ra1 subunit and IL-4Ra, located mainly on B cells, monocytes and dendritic cells[57,58]. Immunofluorescence staining of spleen sections with phosphorylated IL-13Ra1 co-stained with CD19, CD11c and MCPT8 revealed a significant increase of the ratio of pIL-13Ra1+/CD19+ and pIL-13Ra1+/CD11c+ cells upon TBI (CD19: p = 0.0305; CD11c: p = 0.0465; Figure 4D, E and suppl. Figure 3A, B). Interestingly, MCPT8+ basophils also show a significant increase of pIl13-Ra1 positivity (p = 0.0176, Figure 4D, E and suppl. Figure 3C), suggesting a paracrine effect toward B cells and DC but also an autocrine effect.

In fact, the spatial distribution of pIL-13Ra1+ cells in proximity of IL-13+ basophils was assessed, revealing an increased number of pIL-13Ra1+ cells in the vicinity of IL-13+ cells after TBI compared to randomly located coordinates (40µm: p = 0.0029; 60µm: p < 0.0001; 80µm: p < 0.0001; 100µm: p < 0.0001; Figure 4F, G).

This data suggests that basophils trigger IL-13 signaling in adjacent B cells and DCs (as well as on themselves).

### IL-13+ Immune synapses between basophils and nearby immune cells revealed by super-resolution microscopy

Besides paracrine effects, immune cells [59,60] including basophils [61,62] are known to engage in direct physical interaction at so-called immune synapses. Close inspection of confocal immunofluorescence stacks displayed several instances of apparent contact between IL-13+ cells and pIL-13Ra1+ cells, with a significant increase in TBI vs sham controls (p = 0.0035; Figure 5A, B). We sought to elucidate the possibility of IL-13+ immune synapses between basophils and DC or B cells using super-resolution STED microscopy to resolve the IL-13 and pIL-13Ra1 clusters as belonging to distinct cells (in contrast to the hypothesis of them belonging to the same cell, thus being dependent on autocrine signaling). STED imaging was performed using IL-13 and pIL-13Ra1 for the two STED channel and MCPT8 as a basophil marker in the confocal channel; STED imaging revealed the close apposition of one or multiple pIL-13Ra1 clusters with IL-13+ cellular protrusions To ensure that the pIL-13Ra1 clusters belonged to a distinct cell, we plotted the distribution of distance peaks between IL-13 and pIL-13Ra1 clusters; quantification of the distance revealed a larger spread, spanning between 0.5 to 3 µm in the sham samples. Interestingly, for the TBI sections, we observed a large peak at around 500 nm, with an overall spread not superior than 1.5 µm (Figure 5C-G). Cellular migration and motility have been linked to an increase of actin protrusion in the cytoplasm towards the direction of movement[63,64]. We sought to confirm the involvement of actin dynamics in basophils post TBI, by super-resolution STED microscopy. STED imaging was performed using β-actin and pIL-13Ra1 in the two STED channel and MCPT8+ as a basophil marker in the confocal channel. STED imaging revealed increased actin protrusions, in MCPT8+ basophils, towards pIL-13Ra1+ cells in TBI sections compared to sham; quantification of the distance revealed a larger spread, spanning between 0 to 14 micrometers between the two cells in the sham samples. Interestingly, for the TBI sections, we observed a large peak at around 2 micrometres, with an overall spread not superior than 12 micrometers (Figure 5H-L).

**Figure 5:**
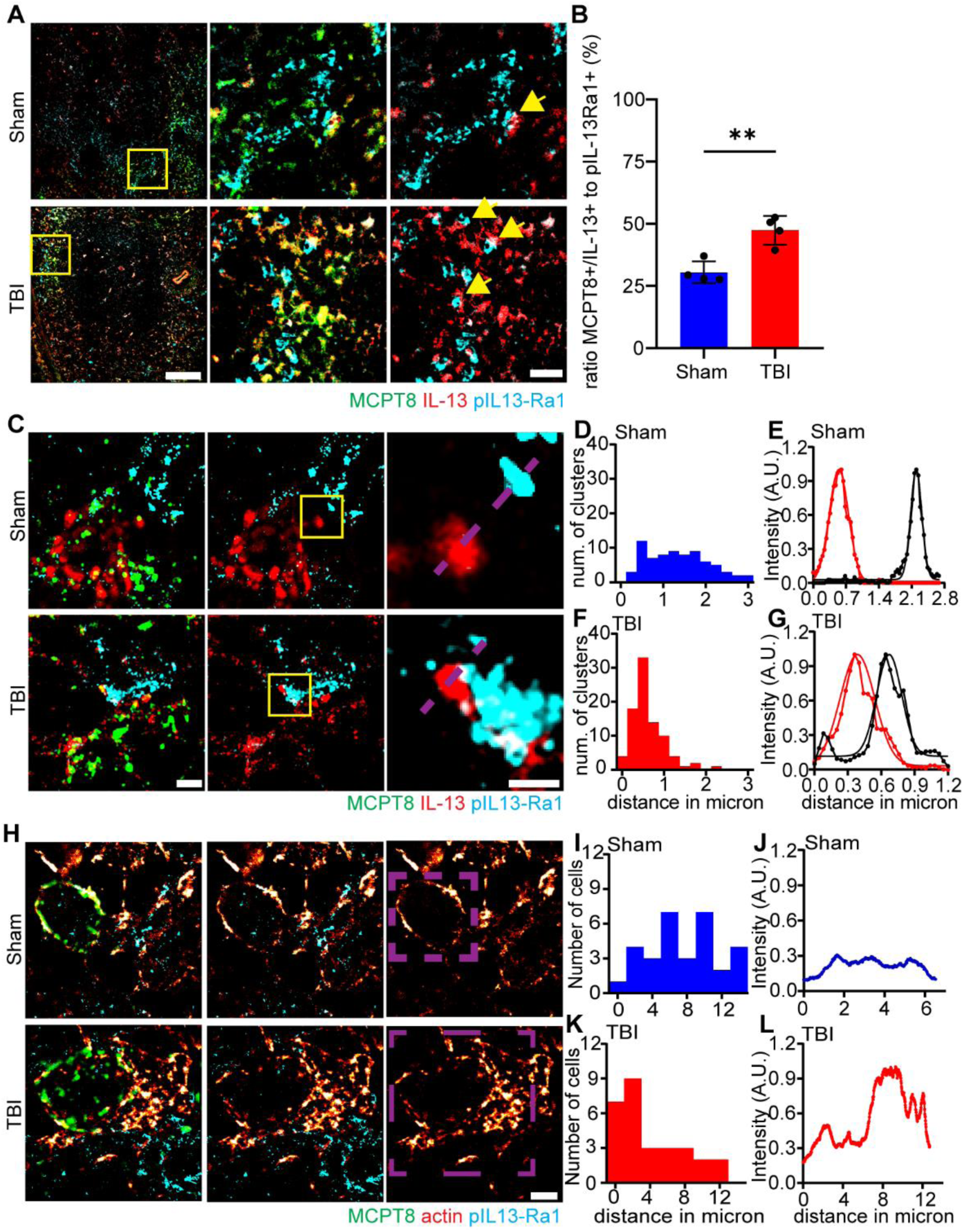
STED imaging reveals IL-13+ Immune synapses between basophils and nearby immune cells. **A, B.** Immunofluorescence staining and confocal imaging showed a significant increase of MCPT8+/IL-13+ cells in close contact with pIL-13Ra+ cells (Sham vs TBI: 31±4% vs 47±6%; p = 0.0035; t = 4.654) post TBI. N = 4, scale bar overview: 100 µm; scale bar insert: 20 µm. **C-G.** STED imaging and intensity plot profiles of spleen sections with MCPT8, IL-13 and pIL-13Ra1 revealed a smaller spread and a closer contact between IL-13 and pIL-13Ra1 clusters post TBI compared to sham. N = 3, scale bar overview: 2 µm; scale bar insert: 1 µm. **H-L.** STED imaging and intensity plot profiles of spleen sections with MCPT8, β-actin and pIL-13Ra1 revealed a smaller spread and a closer contact between β-actin and pIL-13Ra1+ cells post TBI compared to sham. N = 3, scale bar: 2 µm. Data is shown as histogram on frequency distribution to distance and intensity plot profile of representative cell or cluster.

The super-resolution evidence implies an increase in direct contact between basophils and other immune cells upon TBI and the establishment of immune synapses enabling close-range IL-mediated signaling.

### Basophils drive protein-synthesis upregulation in B-cells and DC through IL-13 and long-non-coding RNA NORAD

IL-13 signaling has been reported to be a regulator or mTOR dependent protein synthesis[65–67] and upregulation of metabolic rates and protein synthesis is an early marker of immune cells reactivity[68,69]. Thus, we hypothesized that B-cells and DC receiving basophil-derived IL-13 may respond by upregulating their protein synthesis rate. In fact, we investigated this effect using the phosphorylation of the S6 ribosomal protein as proxy for mTOR activation. Furthermore, we exploited the phosphorylation of 4E-BP1 as a proxy for the enhancement of cap-dependent mRNA translation (by dissociation of eIF4E;[70]). Thin spleen sections were therefore immunostained for pS6 or, in independent experiments, for p-4E-BP1 together with CD19, CD11c and MCPT8. Confirming our hypothesis (and coherent with the phospho-proteomics array, Fig. 1E, F), we detected a significant increase in CD19 and CD11c immunofluorescence for either pS6 or p4E-BP1 immuno-positive cells, after TBI (pS6+/CD19+: p = 0.0210; pS6+/CD11c+: p = 0.0003; p4E-BP1+/CD19+: p = 0.0024; p4E-BP1+/CD11c+: p = 0.0007; Figure 6A-D and suppl. Figure 4A-D). Interestingly, MCPT8+ cells (basophils) displayed only an increased colocalization with pS6, but not with p4E-BP1 immunolabelled cells (Figure 6A, B; Figure 2F-G and suppl. Figure 4E).

**Figure 6:**
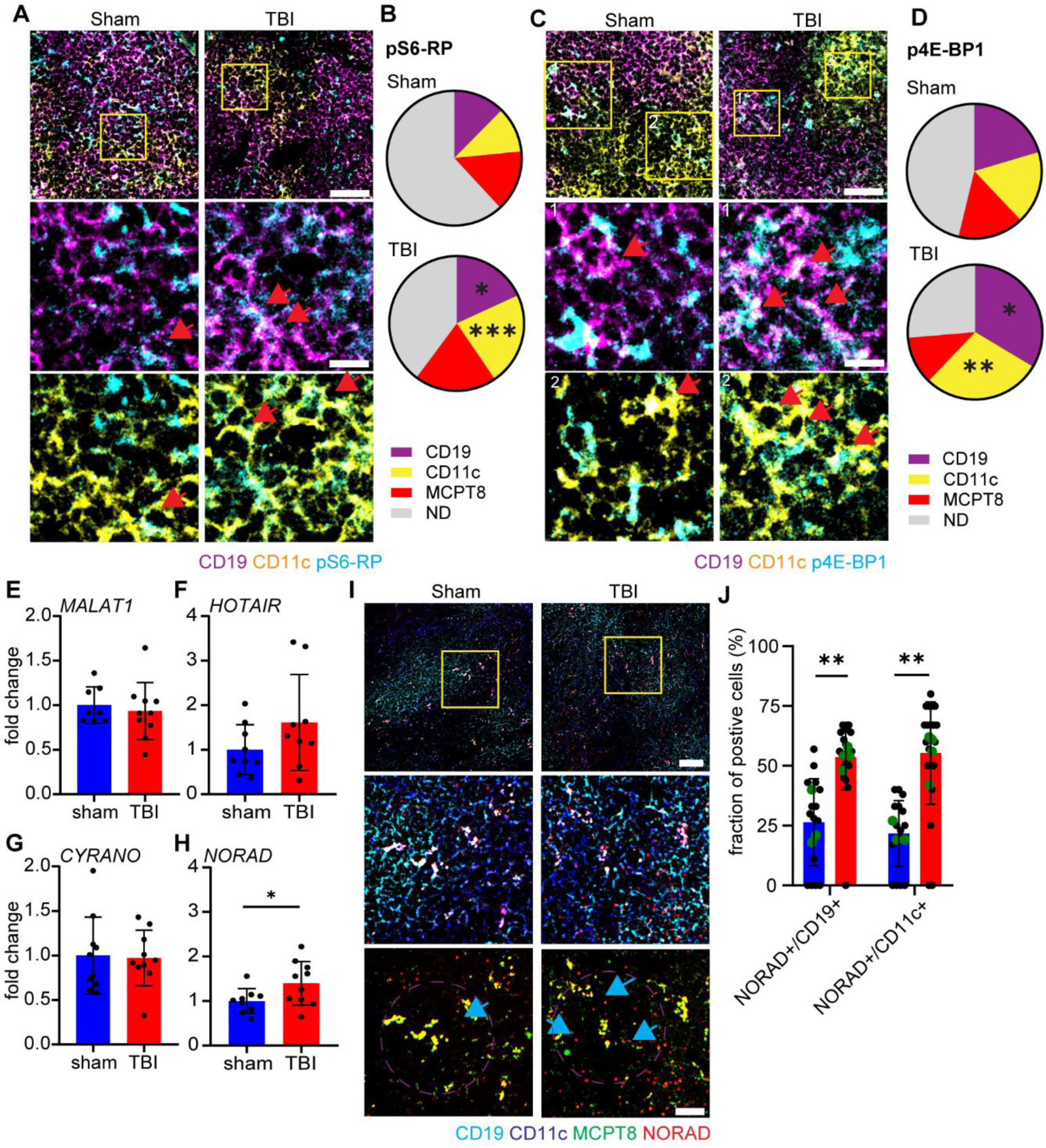
Basophil induces rapid translation response, via IL-13 and NORAD, in B-cells and DCs. **A, B.** Immunofluorescence staining of spleen sections with pS6-RP, CD19 and CD11c showed a significant increase of pS6+/CD19+ (Sham vs TBI: 12±2% vs 18±4%; p = 0.0210; t = 2.864) and pS6+/CD11c+ (Sham vs TBI: 11±2% vs 22±4%; p = 0.0003; t = 6.012) cells post TBI. N = 5, scale bar overview: 50 µm; scale bar insert: 20 µm. **C, D**. Immunofluorescence staining of spleen sections with p4E-BP1, CD19, CD11c and MCPT8 shows a significant increase of p4E-BP1+/CD19+ (Sham vs TBI: 20±5% vs 34±4%; p = 0.0024; t = 4.357) and p4E-BP1+/CD11c+ (Sham vs TBI: 18±4% vs 29±2%; p = 0.0007; t = 5.345) cells post TBI, but no difference in p4E-BP1+/MCPT8+ cells. N = 5, scale bar overview: 50 µm; scale bar insert: 20 µm. **E-H**. Qualitative RT-PCR in spleen samples revealed no differences in lncRNA MALAT1, HOTAIR and CYRANO, but a significant increase in NORAD (Sham vs TBI: 1±0.3 vs 1.4±0.5; p = 0.0484; t = 2.127) post TBI. N = 5. **I, J**. Single lncRNA RNAscope in spleen sections with the probe NORAD and a co-staining with CD19, CD11c and MCPT8 revealed a significant increase in NORAD+/CD19+ (Sham vs TBI: 26±12% vs 54±4%; p = 0.0074; t = 4.341) and NORAD+/CD11c+ (Sham vs TBI: 22±5% vs 55±9%; p = 0.0023; t = 5.701) in close proximity (40 µm) to MCPT8+ cells. Sham N = 3; TBI N = 4, scale bar overview: 50 µm; scale bar insert: 20 µm. Data is shown as pie-charts or mean±SD. *: *p* < 0.05, **: p<0.01, ***: p<0.001.

Protein synthesis upregulation is not only dependent on phosphorylation cascades, but it is also regulated by a number of RNA-dependent mechanisms[71,72]. Among these, the long-non-coding RNA NORAD has been reported to enhance mRNA stability by sequestering PUMILIO proteins, which normally decrease mRNA stability[73,74]. We preliminary screened the possible involvement of NORAD-mediated mechanisms in spleen upon TBI by assessing the overall NORAD expression in whole-spleen extracts, along with 3 lncRNA also involved in immune regulation (namely MALAT1, HOTAIR, CYRANO). Quantitative RT-PCR showed a significant NORAD upregulation in the spleen post TBI (p = 0.0484; Figure 6H) but no difference in the expression of MALAT1, HOTAIR and CYRANO (Figure 6E-G), indicating the specificity of the lcnRNA response.

To provide cellular context to NORAD upregulation, and to verify that NORAD upregulation takes place in B-cells and DCs proximal to basophils, we performed a single-molecule in situ hybridisation for NORAD RNA together with immunohistochemistry for the cell markers CD19, CD11c and MCPT8. The overall density (cells/mm^2^) of NORAD+ cells in the spleen was significantly increased in TBI, with the majority of NORAD+ cells located in the marginal zone of the follicles.

MCPT8+ basophils displayed a strong NORAD signal already in sham samples, which is not changed upon TBI (Suppl. Figure 4F, G). On the other hand, NORAD positivity was assessed by counting the ratio of NORAD+ and either CD19+ or CD11c+ cells in a proximity of 40 micron to an MCPT8+ cell. We showed an increased number of NORAD+/CD19+ and NORAD+/CD11c+ cells in close proximity to MCPT8+ cells compared to their density in proximity of randomly located spleen cells (CD19: p = 0.0074; CD11c: p= 0.0023; Figure 6I, J).

Overall, this data suggests that direct cell-to-cell signaling from basophils to B-cells and DCs results in a rapid protein synthesis and mRNA stability upregulation, through mechanisms involving protein phosphorylation as well as lncNORAD induction.

### TBI induced increase of basophils in the follicles returns back to baseline 3d post injury

To investigate if the basophil recruitment to the splenic follicles persisted beyond the 3h timepoint considered so far, an immunofluorescent staining on thin sections with Ly6G and CCR3 coimmunostained for CD19 was carried out for spleen samples from mice 3 days post TBI (as in Figure 2A). The confocal images revealed no differences in Ly6G+ cells by themselves between sham and TBI (Figure 7A, B), similarly to the 3h time point. Interestingly, we detected no differences in the fraction of CCR3+/Ly6G+ cells post TBI (Figure 7A, C). In line with the number of basophils in the splenic follicles, the expression of CXCL1 mRNA, shown by single molecule in situ hybridisation, was reduced to baseline 3 days post TBI (Figure 7D, E). This suggests that the basophils’ chemoattraction and homing in the spleen persisted for a period of time shorter than 3 days. Thus, immunomodulatory effects of basophils in TBI are short-lived. Pointing towards a reduced chemoattraction of basophils towards the spleen in the subacute phase of TBI.

**Figure 7:**
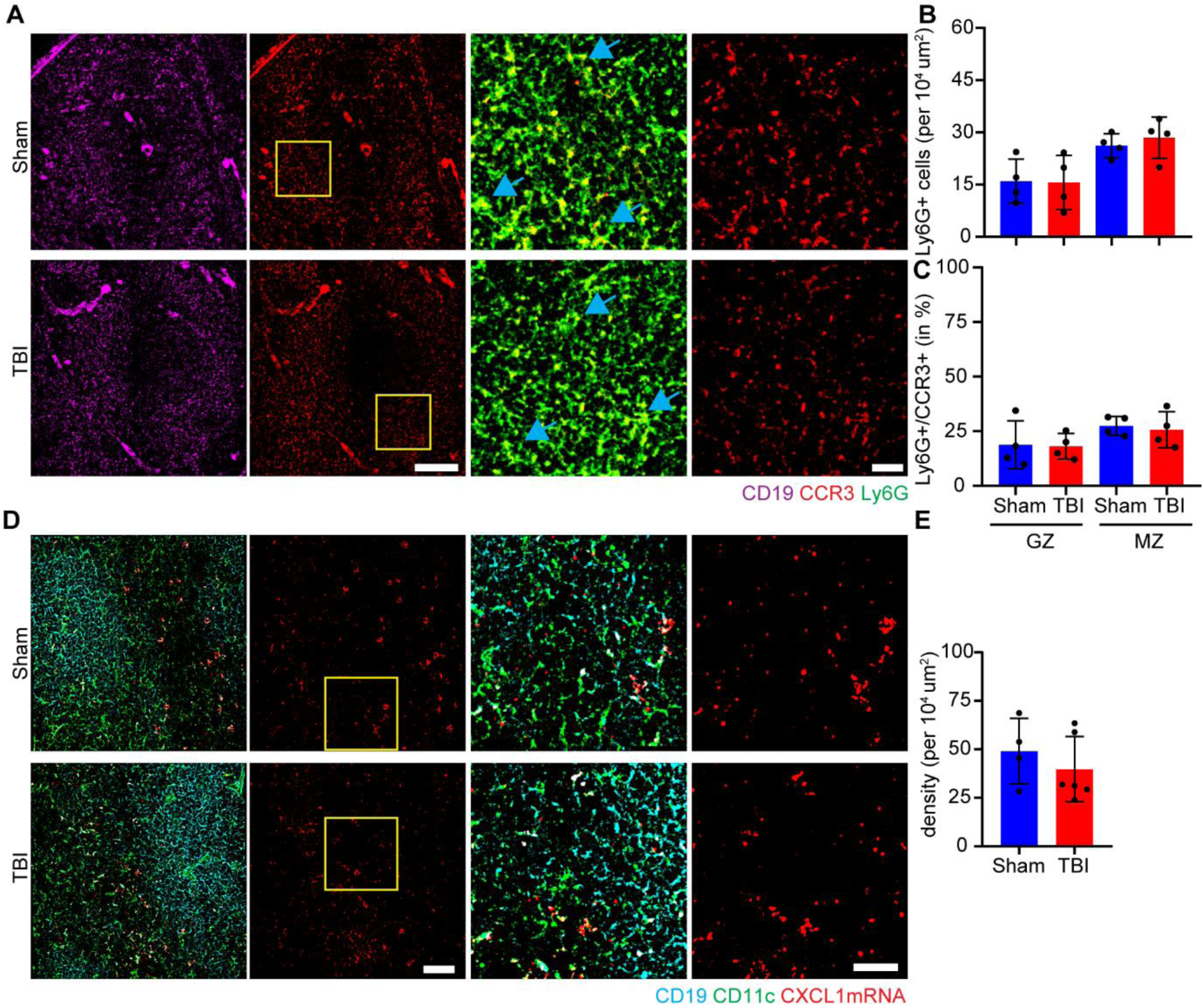
TBI induced increase of basophils returns to baseline 3 days post TBI. **A-C.** Immunofluorescence staining of spleen sections with CD19, Ly6G and CCR3 showed no differences in either the number of Ly6G+ cells nor in Ly6G+/CCR3+ cells 3 days post TBI. N = 4, scale bar overview: 100 µm; scale bar insert: 20 µm. **D, E**. Single mRNA RNAscope revealed no difference in CXCL1 mRNA puncta in the spleen 3 days post TBI. Sham N = 4; TBI N = 6. scale bar overview: 50 µm; scale bar insert: 10 µm. Data is shown as mean±SD.

### Ethanol intoxication prevents the chemotaxis of basophils to the spleen

Ethanol intoxication is a frequent co-morbidity in TBI patients, with 40% of patients showing a positive blood alcohol level upon admission[75] with possible outcome relevance[76–78]. The immunomodulatory effects of ethanol in TBI inflammatory response have been previously investigated in the brain[24,25,79] as well as in the spleen[20,80,81]. Having characterized the basophil recruitment to the spleen as an early event in TBI-associated immune response, here we set out to investigate the effect of ethanol on the basophil phenotype reported. We treated the mice with a single high dose of ethanol (5 g/kg) 30 min before the TBI was administered. Spleen were dissected 3h after the injury and thin sections were immunostained with two sets of staining, on the one hand Ly6G, CCR3 and CD19, and on the other hand pIL-13RA1, CD19, CD11c and MCPT8. The staining revealed no differences in the amount of Ly6G+ cells per area (10^4^ um^2^) in any of the groups (Figure 8A, B). Interestingly, in the case of the density of CCR3+/Ly6G+ cells, ethanol pretreatment prevented the TBI induced upregulation of basophils in the spleen in both the germinal center as well as the marginal zone (Germinal center: p = 0.0093; marginal zone: p = 0.0053; Figure 8A, C). In line with the basophil numbers post TBI with and without ethanol pretreatment, the pIl-13RA1 staining revealed that ethanol prevented the TBI induced pIL-13Ra1 signaling in CD19+, CD11c+ and MCPT8+ cells (CD19: p = 0.0146; CD11c: p = 0.0119; MCPT8; p = 0.0176; Figure 8D, E).

**Figure 8:**
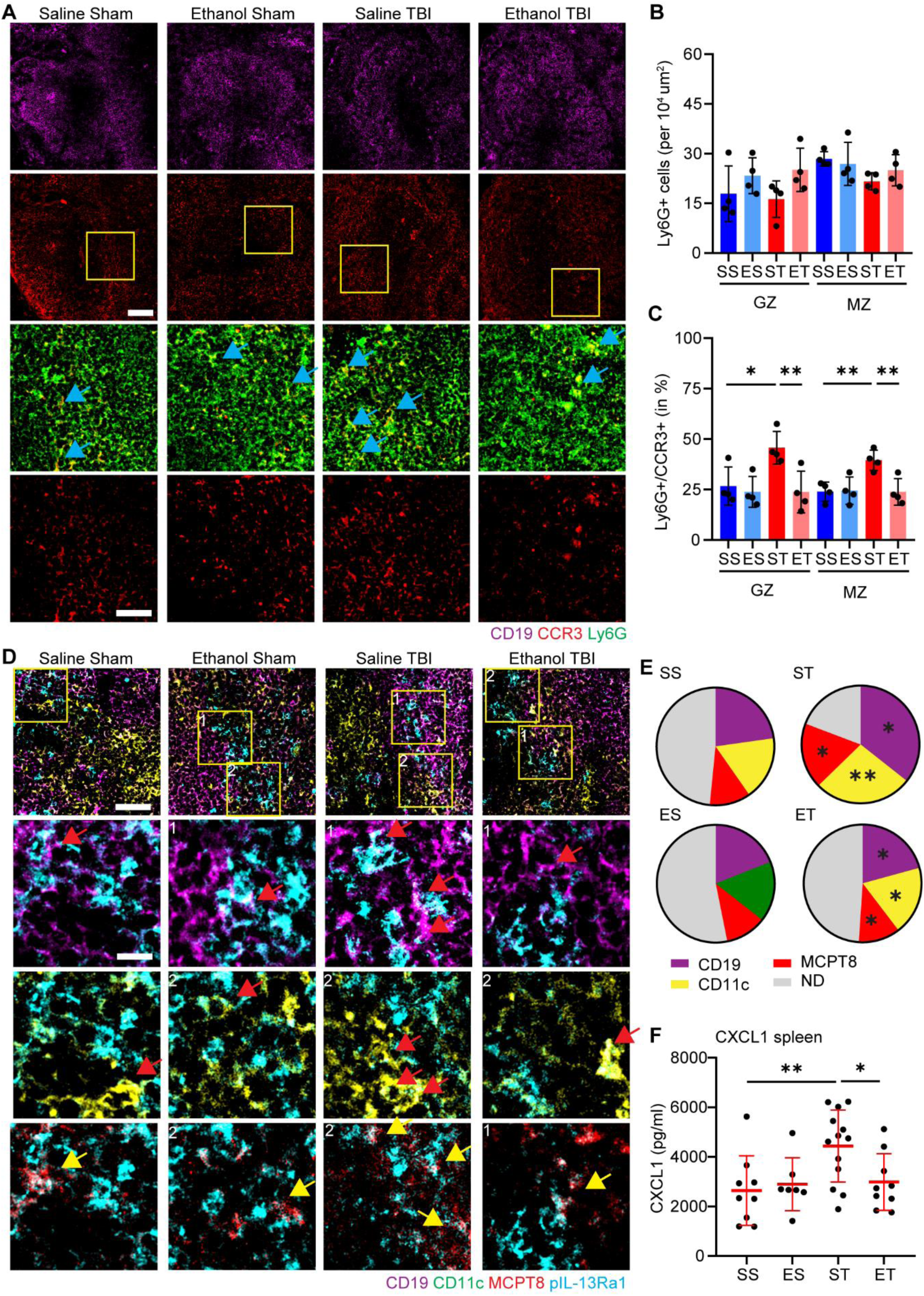
Ethanol pretreatment prevents the chemotaxis of basophils to the spleen. **A-C.** Immunofluorescence staining of spleen sections with CD19, Ly6G and CCR3 show no differences in the number of Ly6G+ cells 3h post TBI (ST) or ethanol-TBI (ET). However, a significant increase was observed in Ly6G+/CCR3+ cells post TBI in the germinal center (SS vs ST: 26.7±9.4% vs45.7±8.1%; p = 0.0219) and marginal zone (SS vs ST: 24±5% vs 40±5%; p = 0.0055) of splenic follicles. With a significant decrease in Ly6G+/CCR3+ cells with ethanol pretreatment, in both the germinal center (ST vs ET: 46±8% vs 24±10%; p = 0.0093) as well as the marginal zone ( ST vs ET: 40±5% vs 24±7%; p = 0.0053). N = 4, scale bar overview: 100 µm; scale bar insert: 20 µm. **D, E.** Immunofluorescence staining of spleen sections with CD19, CD11c, MCPT8 and pIL-13Ra1 showed the significant increase post TBI in pIL-13Ra1+/CD19+ (SS vs ST: 22±4% vs 36±8%; p = 0.0328), pIL-13Ra1+/CD11c+ (SS vs ST: 18±3% vs 27±5%; p = 0.0043) and pIL-13Ra1+/MCPT8+ (SS vs ST: 11±3% vs 18±3%; p = 0.0154). With a significant decrease with ethanol pretreatment, in pIL-13Ra1+/CD19+ (ST vs ET: 36±8% vs 21±7%; p = 0.0146), pIL-13Ra1+/CD11c+ (ST vs ET: 27±5% vs 19±4%; p = 0.0119) and pIL-13Ra1+/MCPT8+ (ST vs ET: 18±3% vs 11±3%; p = 0.0176). N = 4, scale bar overview: 50 µm; scale bar insert: 20 µm. **F.** ELISA assay in a larger cohort of the 4 treatment groups (SS; ES; ST and ET) revealed a significant upregulation of CXCL1 post TBI (SS vs ST: 642±1403 pg/ml vs 4439±1451 pg/ml; p = 0.0066), with a significant decrease with ethanol pretreatment (ST vs ET: 4439±1451 pg/ml vs 2987±1145 pg/ml; p = 0.0303). SS N = 10; ES N = 7; ST N = 13; ET N = 9. Data is shown as mean±SD. *: *p* < 0.05, **: p<0.01.

Furthermore, to verify if the reduced basophil numbers upon ethanol pretreatment is caused by a decreased amount of CXCL1, we performed an CXCL1 ELISA in spleen protein samples. We found a significant downregulation of CXCL1 in the ethanol pretreated samples compared to TBI alone (p = 0.0303; Figure 8F).

These data suggest that recruitment of basophils to the spleen and the subsequent upregulation of IL-13 is prevented by ethanol, further expanding the mechanistic understanding of ethanol immunomodulation.

## Discussion

In the present work we provide evidence for a previously unappreciated mechanism responsible for rapid immunomodulatory response following TBI, namely the rapid recruitment of basophils to spleen lymphoid follicles. Basophils are recruited through chemotactic cues that include CXCL1, deliver an IL-13-mediated signal to B-cells and dendritic cells and contribute to their activation by triggering a protein synthesis response that involve phosphorylation cascades as well as the induction of lncRNA NORAD. The basophils recruitment appears to be transient, possibly because of the downregulation of IL-5 simultaneous to their recruitment, and disappears by 3 dpi. Thus, upon TBI, spleen cells recruit an additional source of IL-13 in the form of the peripheral pool of basophils and this IL13 is delivered to B-cells and DC to shape their early response. Interestingly, this early recruitment of basophils is among the processes altered by ethanol intoxication in TBI.

TBI is marked by systemic immune effects[82], with both a diminished immune response, revealed by reduced levels of monocytes, lymphocytes and natural killer cells[6–8], as well as an increased inflammation, shown by enhanced inflammatory markers in the blood[83,84] and an increase in neutrophils and eosinophils post injury[10,85,86]. Interestingly, in contrast to neutrophils and eosinophils, an increase in basophils post TBI was not found[87]. Which might be due to the overall low numbers (<1%) and short lifespan (2-3 days) of basophils, compared to other granulocytes or leukocytes in general[88,89].

We have shown a significant involvement of basophils in splenic follicles, interacting with B-cells and DCs, using both paracrine mediators as well as cell-to-cell contacts. Basophils have previously been reported to act as very potent antigen presenting cells, inducing both a Th2 response in CD4+ T-cells[61,62,90,91], as well as an CD8+ T-cell response[92], via direct cellular contact or by means of soluble mediators[93]. The interaction and activation of B-lymphocytes by basophils has been reviewed extensively by Merluzzi et al., in 2015[94], focussing on both soluble mediator dependent interaction, as well as cell-to-cell contact between basophils and B-cells. Earlier studies have reported basophils expressing IL-13 to activate B-cells, leading to maturation and induction of IgE[95,96]. More recent reports have also reported the interaction of B-cells with basophils, leading to survival, differentiation and maturation of B-cells into plasma cells in mice[97] and humans[98]. Direct cellular interaction, by immunological synapse, of basophils with dendritic cells has not been reported previously. However, a close cooperation between the two cells, by soluble mediator expression of basophils that activate DCs (e.g. IL-4 and IL-13;[99]) or ROS mediated signaling between basophils and DCs to elicit a Th2 response has been reported[100]. Our data is largely in agreement with previous findings, showing basophils activating B-cells and dendritic cells, with two distinct mechanisms of interaction (paracrine and cell-to-cell contact) as shown by the literature. However, further investigations are needed to disentangle if one cell type is preferred for a specific type of interaction in this context.

What is causing the TBI induced increase of CXCL1 in splenic B-cells and DCs, and what could be the consequences? CXCL1, also known as GRO-a or KC, has been reported as a granulocyte chemoattractant and plays a key role in cancer, by recruiting neutrophils to the tumor environment[101,102]. It has also been reported to play a role in neurological disorders[103,104]. Recent studies identified CXCL1 as a chemoattractant for basophils, by acting on the CXCR2 receptor located on basophils[50]. In addition, several other receptors for CXCL1 have been reported to be expressed on basophils[45], pointing towards the involvement of multiple chemokine receptors for CXCL1 signaling in basophils. The expression of CXCL1 in immune cells is largely induced by an inflammatory insult, either by mediators (cytokines/chemokines;[105,106]), pathogen associated molecular patterns or damage associated molecular patterns (PAMPs and DAMPs;[107,108]). In fact, Burke et al., showed that NF-kB and STAT1 pathways are directly involved in the expression of CXCL1[109]). Furthermore, Hu et al., reported the expression of CXCL1 upon LPS stimulation, and thereby TLR4 signaling, in murine B cells[110]. These reports point towards a TBI induced damage response or inflammatory insult in the spleen, which induces CXCL1 expression in B-cells and DCs. In fact, damage markers are already elevated in the plasma 3h post TBI[111]. Nevertheless, recent studies have shown strong involvement of autonomic innervation in controlling splenic responses via adrenergic and cholinergic signaling[13,17,112]. Furthermore, adrenergic receptor expression in DCs has been reported to be decreased 3h post TBI, which is associated with increased maturation[20]. In fact, electrical induction of sympathetic neurotransmitters decreases expression of CXCL1, IFN-y and IL-6 in the spleen[18]. Thus, a role for the autonomic innervation in contributing to TBI-induced CXCL1 expression may take place along with the effect of systemic damage induced mediators.

IL-13, expressed largely by mast cells, basophils and eosinophils[113], is best known for its T helper 2 (Th2) cell-mediated Type 2 inflammation, resulting in an humoral immune response, promotion of B cell proliferation and induction of antibody production[114–116]. IL-13 has been reported to induce B-cell proliferation and inducing antibody class switching to IgG4 and IgE[94–96]. More recent studies have shown differentiation, maturation and survival of B-cells in response to IL-13, expressed by basophils[97,98]. In fact, when B cells are depleted of the IL-13 Co-receptor, IL-4Ra, a decrease of B-cell responsiveness and IgE expression upon an allergic insult was observed[117]. Furthermore, IL-4Ra has been shown to regulate IgE response in B cells in acute and chronic atopic dermatitis[118]. In the case of dendritic cells, early studies have reported the maturation of DCs upon IL-13, with increased MHC-II and costimulatory factors, like CD40 and CD86[119,120]. With more recent studies showing the Th2 directed immune phenotype of DCs in an IL-13 dependent fashion[121–124]. In fact, inhibition of STAT6, the main transcription factor of IL-13 signaling, was reported to reduce the induction of the Th2 adaptive immunity by reduction of DCs migration[125]. In addition to the IL-13 driven enhanced Th2 response by DCs, mast cell derived IL-13 has been reported to downregulate IL-12 expression in DCs, which in turn inhibits the Th1 cell response[126]. Interestingly, we observed a downregulation of IL-5 following injury, an important activator of basophils[127,128]. Suggesting not only a rapid initial activation of basophils but also a subsequent suppression of their response, as indicated by the decrease in basophil numbers and the concurrent activation of B cells and DCs at 3 dpi. Suggesting a short-lived interaction of basophils with B cells and DCs after TBI. Thus, the rapid activation of B cells and DCs, by basophil derived IL-13, found here may point towards both a differentiation and maturation of follicular B-cells as well as a maturation of DCs in the spleen upon TBI, followed by a potential Th2 adaptive immune response.

This study revealed a TBI induced basophil to adaptive immunity crosstalk, with an enhanced translational response in B-cells and DCs driven by phosphorylation of S6 and 4E-BP1 and an increase transcription of the lncRNA NORAD, which has been reported to induce a rapid translation by enhancing mRNA stability[73,74]. The enhanced translational response was driven by IL-13 signaling, which has been reported to induce mTOR dependent protein synthesis[65–67]. The enhanced metabolic acitivtiy of B cells and DCs, induced by basophils, may point toward an increase in Th2 cell development, as described previously[91,114,129,130]. In fact, the systemic consequences post TBI has been associated with a shift in Th polarization, showing a decrease in Th1 response and an increase in Th2 response[131–134], which may be a further risk for sepsis, multi-organ failure and systemic inflammatory response syndrome following injury[135]. In contrast, an increase in the proinflammatory cytokine TNF-α upon DC maturation post TBI has been reported[20], which might point to an increase of the Th1 response[136,137]. However, the involvement of specific DC subsets might reveal the Th directed reactivity in the spleen post TBI. In fact, CD8a+ and CD8a-DC subsets have been reported to induce a Th1 and Th2 response respectively[138–140]. The involvement of a specific Th response in the spleen post TBI remains elusive and requires further in-depth investigations.

The present work displays a few limitations. First, the phosphoproteomic screening reveals multiple signaling cascades, only some of which are traced to basophils, because of their broad expression across multiple immune cells subpopulations (e.g., interferon response, tnf signaling). Thus, additional processes are unfolding in the splenic lymphoid tissue for which it is not possible to provide a precise cellular deconvolution. Second, the identification of the subpopulations responding to the basophil-derived IL-13 is influenced by the sensitivity of the immunofluorescence assay; it is conceivable that once released in the extracellular milieu, more populations (including T-cells) may respond to this cytokine, but their protein-synthesis induction is not detectable with current methods. Thus, the impact of basophils recruitment may extend beyond B-cells and DC. Third, the focus on early events occurring within the preserved architecture of the splenic lymphoid tissue interfered with the follow-up of the same cells at later stages. As such, the ultimate outcome of IL-13 signaling can only be speculated, based on the known effects of IL-13 on antigen presentation of DC[119,120] and on B-cells proliferation[94–96].

In conclusion, here we have demonstrated that TBI results in a coordinated immunomodulatory effort in the lymphoid tissue of the spleen, which involve the active recruitment of granulocyte subpopulations, such as basophils, to deliver critical cytokine signaling to DC and B cells, possibly shaping immune responses to the brain injury. Thus, the basophil-IL-13 axis in TBI provides a new window of opportunity to manipulate acute and long-term immune reactions to TBI for therapeutic purposes.

## Acknowledgements

We thank all the members of the CRC 1149 for their scientific input and discussion. We would like to thank Prof. Anita Ignatius for the access to the histology facility. Technical support by Thomas Lenk was highly appreciated.

## Authors contributions

FoH and FR designed and conceived the study. FoH, JZ and FS performed antibody array assays and ELISA assays. SK performed bioinformatic analysis of the arrays. FoH and GY performed Immunofluorescence staining and single mRNA in situ hybridisation. FoH performed qPCR, confocal, super-resolution imaging and image analysis. FR, MH-L, MP and FoH cooperated in the critical analysis of the data and the preparation of the initial draft. FR, MH-L, MS and FoH contributed to the final version of the manuscript. All authors read and approved the final manuscript.

