## supplementary files for "Basophils activate splenic B cells and Dendritic cells via IL-13 signaling in acute Traumatic Brain Injury"

**Supplementary Table 1: List of antibodies and lncRNA sequences.**

| Antigen (primary antibody) | Catalogue number | Company | Dilution |
|---|---|---|---|
| Guinea pig anti CD11c | HS-375 004 | Synaptic Systems | 1:500 (IF); 1:250 (ISH) |
| Rabbit anti CD19 | ab245235 | Abcam | 1:200 (IF); 1:100 (ISH) |
| Mouse anti CD19 | 14-0199-82 | Invitrogen | 1:100 |
| Rat anti CD4 | 14-9766-82 | E-bioscience | 1:100 (IF); 1:50 (ISH) |
| Rat anti Ly6G | 127632 | Biolegend | 1:300 |
| Rabbit anti CCR3 | NB100-56335 | Novus | 1:100 |
| Rat anti MCPT8 | 647402 | Biolegend | 1:200 (IF); 1:100 (ISH) |
| Rabbit anti S6-RP (phosphor) | 2211S | CST | 1:200 (IF); 1:1000 (WB) |
| Mouse anti IL-13 | SC-393365 | Santa Cruz | 1:50 |
| Rabbit anti IL-13Ra1 (phosphor) | PA5-38607 | Invitrogen | 1:50 |
| Rabbit anti 4E-BP1 (phosphor) | 9455P | CST | 1:500 |
| Rabbit anti ERK1/2 (phosphor) | 9101S | CST | 1:1000 (WB) |
| Mouse anti beta-actin | 69009-I-Ig | Proteintech | 1:500 |
| **Antigen (secondary antibody)** | **Catalogue number** | **Company** | **Dilution** |
| Donkey anti guinea pig 405 | ab175678 | Abcam | 1:500 |
| Goat anti guinea pig 488 | A11073 | Invitrogen | 1:500 |
| Donkey anti mouse 405 | ab175658 | Abcam | 1:500 |
| Donkey anti mouse 568 | A10037 | Invitrogen | 1:500 |
| Donkey anti mouse 647 | A31571 | Invitrogen | 1:500 |
| Donkey anti rabbit 647 | A21207 | Invitrogen | 1:500 |
| Donkey anti rabbit 568 | A10042 | Invitrogen | 1:500 |
| Donkey anti rat 488 | A21208 | Invitrogen | 1:500 |
| FluoTag-X2 anti mouse AbberriorStar 580 | N1202-Ab580-S | Nanotag | 1:500 |
| FluoTag-X4 anti Rabbit AbberiorStar 635p | N2404-Ab635P-S | Nanotag | 1:500 |

| Gene | Sequence |
|---|---|
| MALAT1 | forward: 5’ -atagcccaggaaagagtgcg- 3’<br>reverse: 5’ - gcttcaccaccacatccgta- 3’ |
| HOTAIR | forward: 5’ -gcgccaacgtagaccaaaag- 3’<br>reverse: 5’ -taccgatgttggggacctct- 3’ |
| CYRANO | forward: 5‘ -aggttctgtggcgtgagttg- 3‘<br>reverse: 5‘ -actgcggtcaactgtgctta- 3‘ |
| NORAD | forward: 5’ -tcctgagttgaccgcattgt- 3’<br>reverse: 5’ -ctttccactcacggaccaca- 3’ |

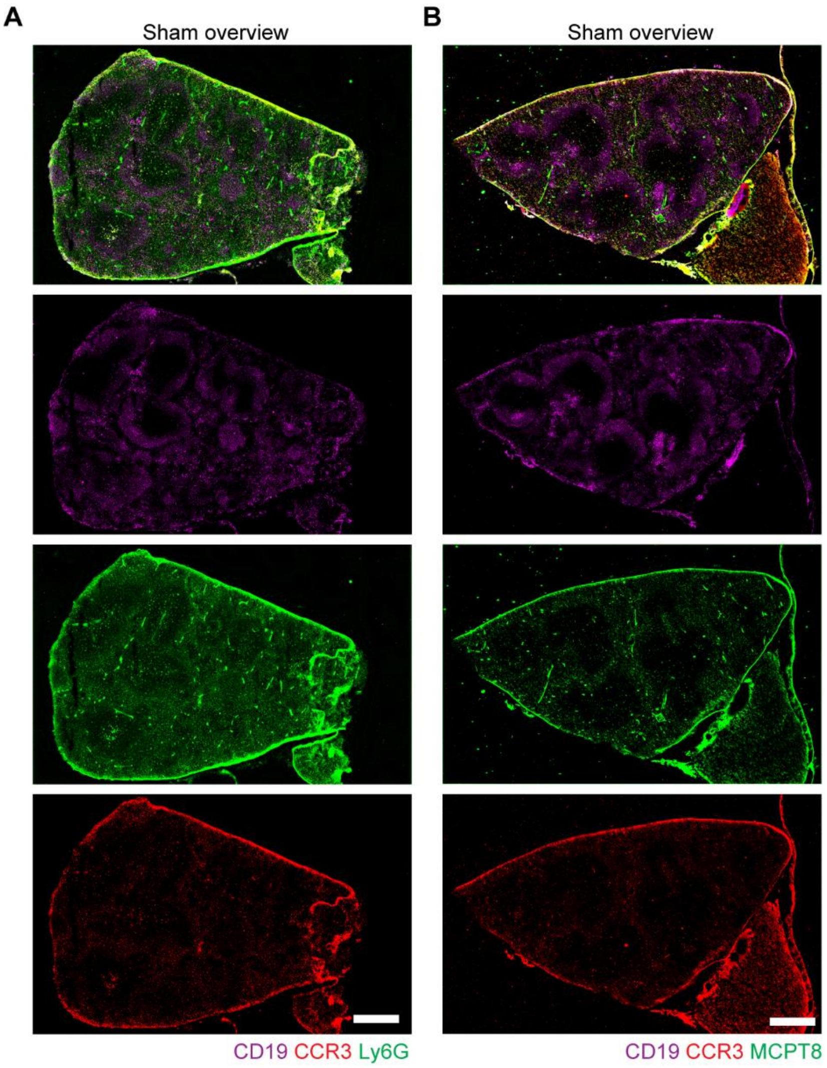

**Supplementary Figure 1: Overview images of thin spleen sections.**

**A.** Immunofluorescence staining and imaging of CD19, Ly6G and CCR3 in thin spleen sections showed the dishomogeneous distribution of CD19 and CCR3, and homogeneous distribution of Ly6G. CD19 and CCR3 showed main localization in the white pulp, marginal zone and follicles, Ly6G showed a localization in both the white and red pulp. **B.** Immunofluorescence staining and imaging of CD19, MCPT8 and CCR3 in thin spleen sections showed the dishomogeneous distribution of CD19, MCPT8 and CCR3. With a main localization in the white pulp, marginal zone and follicles. Scale bar 500 µm.

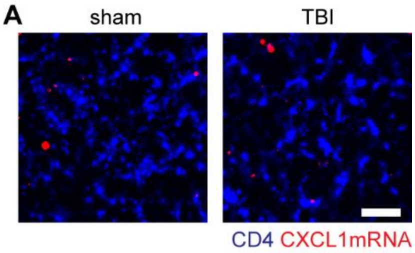

**Supplementary Figure 2: No expression of CXCL1 in CD4+ T-Cells.**

**A.** Single mRNA in situ hybridization of CXCL1 co-stained with CD4 showed no expression of CXCL1 in CD4+ cells in either sham nor TBI. N = 5. scale bar: 10 µm.

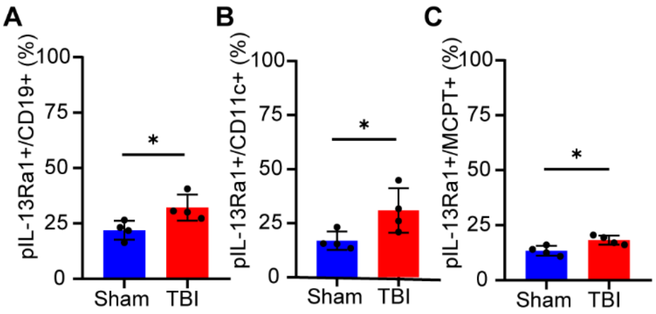

**Supplementary Figure 3: Basophils activate B-cells and DCs via IL-13Ra1 phosphorylation.**

**A-C.** Analysis of the immunofluorescence staining of spleen sections with CD19, CD11c, MCPT8 and pIL-13Ra1, depicted in barplots. Corresponding to Figure 4D. N = 4. Data is shown as mean±SD. *: $p < 0.05$.

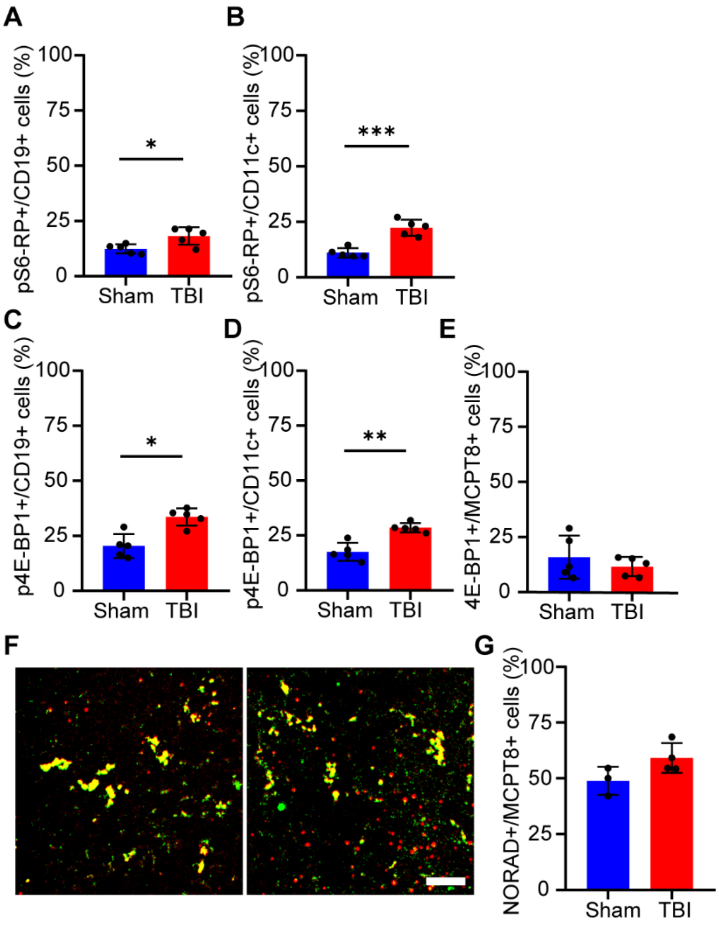

**Supplementary Figure 4: Basophils induce fast translational response in B-cells and DCs post TBI.**

**A, B.** Analysis of the immunofluorescence staining of spleen sections with CD19, CD11c and pS6-RP, depicted in barplots. Corresponding to the representative images in Figure 6A. N = 5. **C-E**. Analysis of the immunofluorescence staining of spleen sections with CD19, CD11c, MCPT8 and p4E-BP1, depicted in barplots. Corresponding to the representative images in Figure 6C. N = 5. **F, G**. Single lncRNA RNAscope in spleen sections with the probe NORAD and a co-staining with MCPT8 revealed a high amount of NORAD in MCPT8+ cells, with no significant differences between sham and TBI (Sham vs TBI: 49±6% vs 59±7%). Sham N = 3; TBI N = 4; scale bar insert: 20 µm. Data is shown as mean±SD. *: $p < 0.05$, **: p<0.01.

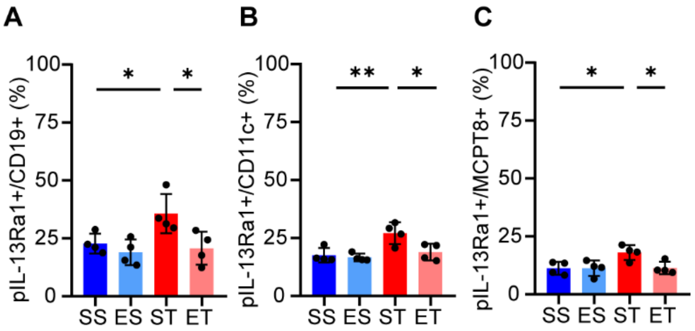

**Supplementary Figure 5: Ethanol pretreatment prevents IL-13Ra1 phosphorylation in basophils, B-cells and DCs.**

**A-C.** Analysis of the immunofluorescence staining of spleen sections with CD19, CD11c, MCPT8 and pIL-13Ra1, depicted in barplots. Corresponding to the representative images in Figure 8D. N = 4. Data is shown as mean±SD. *: $p < 0.05$, **: p<0.01.
